# Optogenetic Instruction of Cell Fate by Temporal Patterning of Mechanobiological Signals

**DOI:** 10.1101/2022.09.14.508003

**Authors:** Rocío G. Sampayo, Mason Sakamoto, Sanjay Kumar, David V. Schaffer

## Abstract

During the intricate process by which cells give rise to tissues, embryonic and adult stem cells are exposed to diverse mechanical signals from the extracellular matrix (ECM) that influence their fate. Cells can sense these cues in part through dynamic generation of protrusions, modulated and controlled by cyclic activation of Rho GTPases. However, it remains unclear how extracellular mechanical signals regulate Rho GTPase activation dynamics and how such rapid, transient activation dynamics are integrated to yield long-term, irreversible cell fate decisions. Here, we report that ECM stiffness cues alter not only the magnitude but also the temporal frequency of RhoA and Cdc42 activation in adult neural stem cells (NSCs). Using optogenetics to control the frequency of RhoA and Cdc42 activation, we further demonstrate that these dynamics are functionally significant, where high- or low-frequency activation of RhoA and Cdc42 drives astrocytic or neuronal differentiation, respectively. In addition, high-frequency Rho GTPase activation induces sustained phosphorylation of the TGFβ pathway effector SMAD1, which in turn promotes astrocytic differentiation. By contrast, under low-frequency Rho GTPase stimulation, cells fail to accumulate SMAD1 phosphorylation and undergo neurogenesis. Our findings reveal the temporal patterning of Rho GTPase signaling and the resulting accumulation of a SMAD1 signal as a critical mechanism through which ECM stiffness cues regulate NSC fate.

## Main

From early embryogenesis and into adulthood^1, 2^, the concerted action of stem cell division and differentiation builds and maintains tissue homeostasis. Differentiation of stem cells into specific lineages is orchestrated by complex biochemical and biophysical signals that vary temporally and spatially^3–5^. For example, mechanical signals in stem cell niches can fluctuate due to transient changes in cell density, induced by cell proliferation and migration, as well as the dynamic deposition of specific ECM molecules by these cells^6^. In addition, it has been proposed that cells sense and decode ECM mechanical cues into intracellular signals by generating membrane protrusions in the order of seconds to minutes, exerting force and deforming underlying physical structures, processes by which cells can repeatedly and locally probe the viscoelastic properties of their environment^4, 7^. Moreover, substrate stiffness has been shown to determine the morphology and dynamics of these exploratory membrane protrusions in several cell types^4, 7–11^. However, whether static substrate stiffnesses affect these dynamic cytoskeletal processes and accompanying activation of signals such as Rho GTPases in stem cells, and whether such fast, reversible mechanisms can be integrated into long-term, irreversible effects on cell fate commitment, are still largely unknown.

The waves of exploratory protrusions are thought to arise due to non-specific adhesion mechanisms^10, 11^, triggering downstream contraction/relaxation events of the acto-myosin cytoskeleton that are controlled by Rho family of GTPases^12^. The reversible nature of Rho GTPases, which switch between GTP-bound active and GDP-bound inactive states^13^, enables cells to sustain cycles of extension and retraction of protrusions, critical for mechanosensing as well as directional migration. These GTPases play critical roles in controlling rapid cytoskeletal network effects as well as long-term signal transduction events that determine gene expression and ultimately determine cell fate^4, 14^.

We previously showed that ECM stiffness instructs cell fate in adult hippocampal neural stem cell (NSC) through a mechanism involving the Rho GTPases RhoA and Cdc42^3, 15, 16^. These NSCs reside in the subgranular zone (SGZ) of the hippocampus and give rise to neurons and glia that play critical roles in learning, memory, and pattern recognition^17, 18^. SGZ NSCs live at the interface between two hippocampal layers with sharply different stiffnesses, which likely exposes them to diverse mechanical inputs^19, 20^. We showed in vitro that NSCs exhibit mechanical memory, wherein they are sensitive to the elastic properties of the substrate between the first 12 to 36 hours of differentiation in culture, after which they are irreversibly committed to a certain fate that arises 4-6 days later^15, 16^. Other stem cell types such as mesenchymal stem cells (MSCs) also exhibit mechanical memory, and Rho GTPase activity has also been shown to be critical for lineage commitment^21, 22^. However, it remains unknown whether static ECM stiffnesses produce static or dynamic activation of Rho GTPases, and whether dynamic activation could be integrated into long-term, persistent signal transduction events necessary to irreversibly determine cell fate beyond the temporal window of mechanosensitivity.

Using NSCs as a model for adult multipotent stem cells, we show that ECM stiffness determines not only the magnitude but, strikingly, also the frequency of Rho GTPase oscillatory activation, with stiffer substrates promoting higher activation frequencies. We then employed optogenetically-controlled RhoA and Cdc42 variants we previously developed^23^ to investigate how this mechanosensitive oscillatory activation of Rho GTPases influences NSC fate decisions. Combining this technology with single-cell differentiation profiling, we reveal that NSC fate commitment is determined by the frequency of oscillatory activation of RhoA and Cdc42 during a critical temporal window. High-frequency activation resulted in persistent actin cytoskeleton subcellular reorganization and downstream phosphorylation and nuclear translocation of SMAD1/5, a transcription factor that promotes astrocytic differentiation. By contrast, low-frequency activation yielded only intermittent activation of SMAD1/5 that was insufficient to promote astrocytic fates and instead favored neuronal differentiation. ECM stiffness thus determines the frequency of Rho GTPases activation, and these dynamic signaling inputs are temporally integrated, through actin cytoskeleton and SMAD1/5 dynamics, into irreversible stem cell fate decisions.

## Results

### Extracellular matrix stiffness controls the frequency of oscillatory RhoA activation in adult neural stem cells

To investigate how ECM stiffness regulates RhoA activation in adult multipotent stem cells, we engineered adult hippocampal NSCs to express a previously described^24^ fluorescence resonance energy transfer (FRET)-based RhoA biosensor that enables quantification of RhoA activity via live-cell imaging (Fig. 1a). We used polyacrylamide (PA) hydrogels that are tunable over a broad range of stiffnesses encompassing the stiffness range of brain tissue and covalently conjugated the surfaces with laminin^25^. We then measured RhoA activation dynamics during the critical temporal window of mechanosensitivity we previously identified for these cells (12-36 hours after induction of differentiation on PA gels)^16^ in the presence of biochemical cues that over 7 days induce mixed differentiation into primarily neurons and astrocytes^15^ (Fig. 1b). Ratiometric FRET analysis revealed that cells on either soft (520 Pa) or stiff (73 kPa) PA hydrogels exhibited heterogeneous, localized subcellular regions with high (average normalized FRET index amplitude ≥ 0.4) or low RhoA activation (average normalized FRET index amplitude ≤ 0.2). Stiff substrates exhibited more abundant regions with high RhoA activity, whereas soft substrates showed more abundant regions with low activity, consistent with our previous ELISA-based GTPase activation assay measurements indicating ∼1.8 fold higher activity of RhoA on stiff substrates^15^ (Fig 1c and Supplementary Fig. 1a). However, temporal analysis of RhoA activation within different regions (approximately ∼400 nm in diameter [Fig. 1d]), strikingly revealed an oscillatory pattern in RhoA activation with significantly different frequencies on soft vs. stiff substrates (Fig. 1c, e and f). That is, NSCs seeded on soft substrates had peaks of RhoA activation every ∼45-60 minutes, whereas RhoA activation peaked every 5-10 minutes on stiff substrates (Fig. 1d-f). RhoA activation thus spontaneously oscillates in NSCs at a frequency modulated by the elastic properties of the substrate (Fig. 1g).

**Figure 1.**
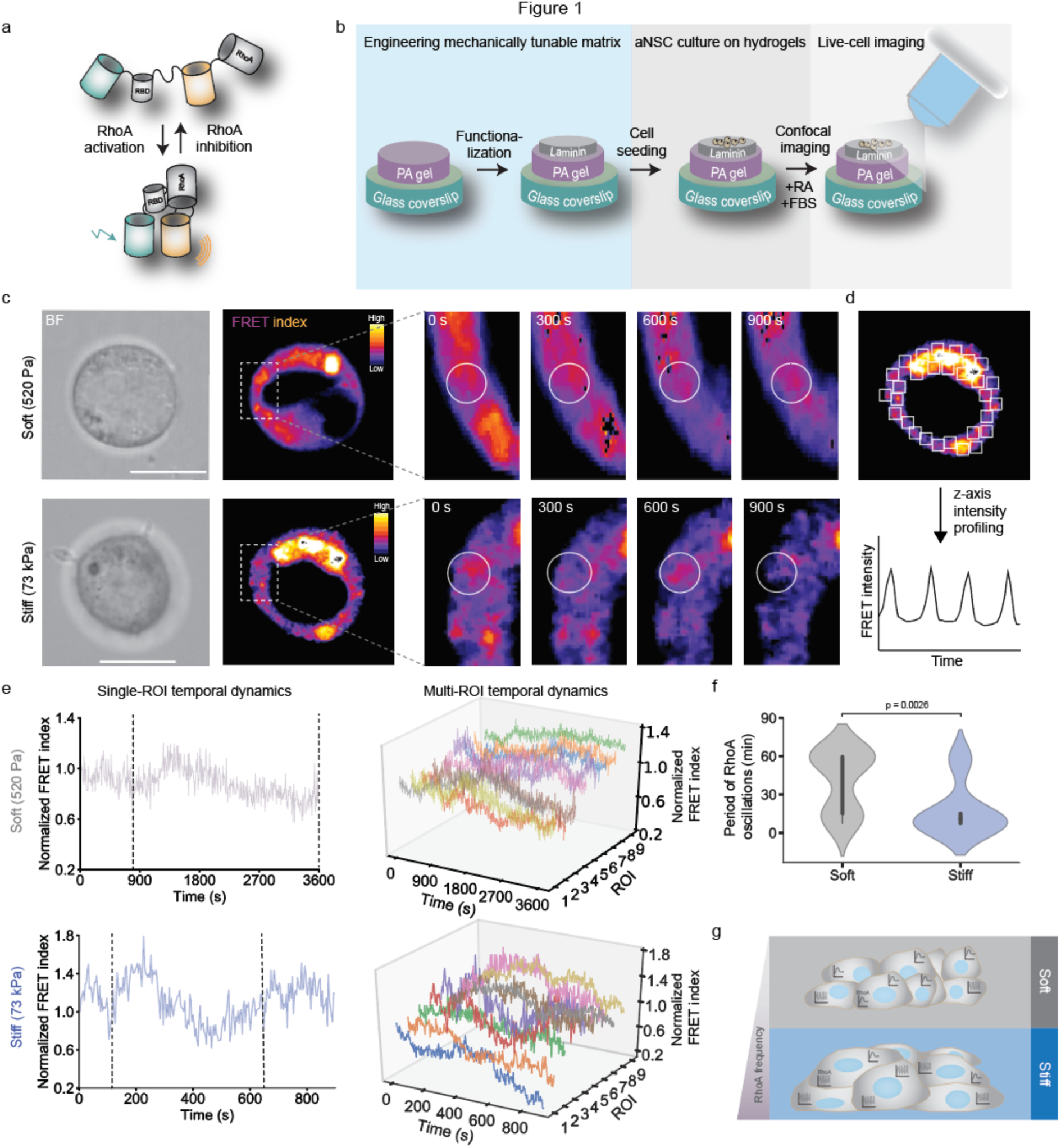
Frequency of RhoA activation is modulated by substrate elasticity in NSCs. **a**, Representation of the FRET biosensor used. **b,** Schematic representation of the PA hydrogel model used of tunable stiffness **c,** Representative images of temporal dynamics of RhoA activation measured by a single-chain FRET biosensor in NSCs seeded on PA hydrogels of different stiffnesses. High FRET index represents high degree of activation of RhoA. Scale bar 10μm**. d,** FRET index temporal profile was tracked in a total of 20 ROIs per cell (400nm^2^ squares); 10 cells per condition were analyzed. **e,** Temporal profile of FRET index in cells seeded on soft or stiff substrates. Left column: representative temporal profile for single-ROI dynamics. Right column: plotted are representative FRET temporal dynamics across 10 ROIs in all cells analyzed. FRET index at each timepoint was normalized to *t*=0. Temporal FRET index profiles were smoothed and detrended to filter out monoexponential decays due to photobleaching, and peaks were counted and averaged to calculate oscillation frequencies. **f,** Violin plots showing RhoA activation frequency distribution between soft and stiff. Student’s t-test was used to evaluate statistical significance. **g,** Cartoon depicting stiffness-dependent RhoA oscillatory activation in NSCs.

We further assessed whether protrusion frequency, which we would expect to be closely connected to RhoA activation, may be sensitive to substrate elasticity. We found that on stiff substrates cells exhibit a pattern of oscillatory neurite extension with a period of 6 min, comparable to the 5-10 min gap between peaks of RhoA activation when cells are seeded on these substrates (Supplementary Fig 1b and c). On soft substrates, we were not able to detect neurite re-extension after retraction (or neurite retraction after extension) within the temporal window analyzed, suggesting that if there is an oscillatory pattern of neurite extension on soft substrates, the period would be longer than the analyzed timeframe. These data suggest that NSCs exhibit oscillatory patterns of neurite extension and retraction, and that the frequency of these oscillations is modulated by substrate stiffness. These fast, spontaneous bursts of neurite extension/retraction found on stiff substrates may contribute to high-frequency oscillatory RhoA activation in NSCs.

### Optogenetic RhoA and Cdc42 enable high spatiotemporal resolution control of GTPase activation in NSCs

Having found that ECM stiffness modulates not only the average magnitude but also the frequency of RhoA activation, we wondered whether Rho GTPase oscillation frequency could impact NSC fate commitment. Standard strategies to manipulate Rho GTPase activation such as biochemical activators of Rho GTPase pathways (LPA or calpeptin) lack temporal resolution due to cellular diffusion constraints. Moreover, genetic perturbation of Rho GTPases typically involves expression of constitutively active (CA) or dominant negative (DN) mutants, which also lack rapid temporal tunability. To overcome these limitations, we previously placed both RhoA and Cdc42 under optogenetic control^26^, enabling Rho GTPase activation with high spatiotemporal resolution. In this system, full length RhoA or Cdc42 was fused to a truncated *Arabidopsis Thaliana* Cryptochrome 2 (Cry2), hereafter named “optoRhoA” and “optoCdc42”, such that blue-light illumination induces clustering of Cry2 and the activation of the accompanying Rho GTPase^26^ (Fig. 2a). This work complemented the previously reported optogenetic Rac1, which used a different strategy^27^.

**Figure 2.**
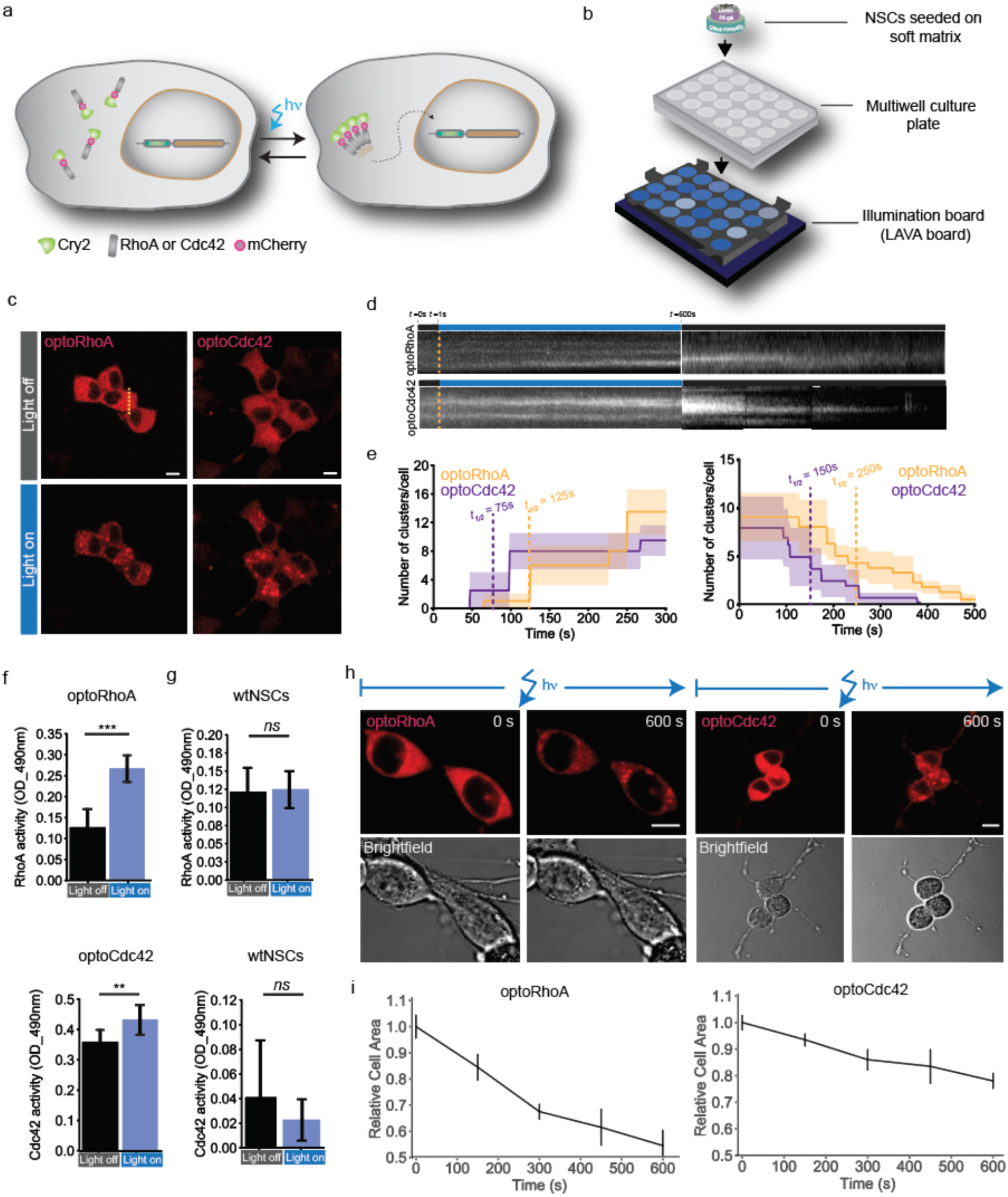
Optogenetic RhoA and Cdc42 for optical control of GTPases dynamic activation in NSCs. **a**, Schematic of optogenetic RhoA/Cdc42 pathway activation (‘optoRhoA’ or ‘optoCdc42’) in NSCs. **b,** Schematic of LAVA illumination board utilized for optogenetic stimulation. **c,** Live-cell imaging of mCherry-optoRhoA or mCherry-optoCdc42 at *t*=0 (top images) or after 15 minutes of blue light stimulation (bottom images). Scale bar 10μm **d,** Kymographs of mCherry-optoRhoA or mCherry-optoCdc42 corresponding to the dashed line. **e,** Representative plot of single-cell cluster formation and dissociation over time. Similar behavior was observed in all cells for which cluster formation was measured. *T*_1/2_, time at which a half-maximal number of visible clusters was detected. **f,** Rho and Cdc42 activity were measured by ELISA-based GTPase activation assays (G-LISA) after 10 minutes of blue light stimulation at 0.5 mW/mm^2^ of optoRhoA or optoCdc42 NSCs seeded on soft substrates. Data are represented as mean ± S.D (n=6 biological replicates) **g,** Rho and Cdc42 was measured by G-LISA in wtNSCs after 10 minutes of blue light stimulation at 0.5 μW/mm^2^. Data are represented as mean ± S.D (n=6 biological replicates). **h,** Representative time-lapse images of optoRhoA and optoCdc42 NSCs upon 10 minutes of opto-stimulation. Contraction was determined with by measuring cell area over time with the software Fiji^60^. Resulting cell areas at each timepoint were plotted relative to *t*=0. Scale bar 10μm**. i**, Quantification of h. Data are represented as mean ± S.D (n=50 cells per condition). \*\*\**P* < 0.005, \*\**P* < 0.01, \**P* < 0.05. ns, not significant.

To precisely control the amplitude and frequency of illumination for optostimulation, we used our recently reported, programmable photostimulation devices for light activation at variable amplitudes (“LAVA” boards)^28, 29^ (Fig. 2b). These devices allow for controlled illumination of multiwell plates in routine cell culture, at user-defined frequencies (10 ms resolution) and amplitudes (0.005 μWmm^2^ resolution) that are specified independently for each well. Cells were seeded on soft (520 Pa) PA hydrogels (Fig. 2b), which provide the lowest average activation background for RhoA and Cdc42 (Fig. 1c)^15^. Blue light stimulation induced clustering and plasma and perinuclear and membrane localization of these proteins, coinciding with previous reports of active Rho GTPase subcellular distribution^13, 26^. Clustering association (t_1/2_ Cdc42 = 75 s, t_1/2_ RhoA = 125 s) and dissociation kinetics (t_1/2_ Cdc42 = 150 s, t_1/2_ RhoA = 250 s) were consistent with our prior work in fibroblasts^26^ (Fig. 2c-e) and in a temporal range useful for simulating endogenous Rho GTPase fluctuations seen in NSCs (Fig. 1).

Furthermore, we measured how strongly opto-stimulation activated RhoA and Cdc42 activation using ELISA-based GTPase activation assays, which showed a nearly 2-fold activation of RhoA and ∼1.3 fold activation of Cdc42 upon static illumination (Fig. 2f), in contrast to wild-type NSCs (wtNSCs) in which no elevation was observed upon illumination (Fig. 2g). To further examine the functional consequences of optogenetic RhoA or Cdc42 activation, we evaluated morphological effects. A short (10-minute) pulse of light was sufficient to trigger fast cellular contraction in optoRhoA cells (Fig. 2h and i), with milder cellular contraction with optoCdc42 (Fig. 2h and i). wtNSCs did not show morphological effects when exposed to blue light (Supplementary Fig. 2a). These results validate the use of this light-induced oligomerization system for rapid activation of RhoA and Cdc42, which increased cellular contractility as described for these proteins in NSCs^15^ and other cell types^30, 31^.

### Oscillatory Rho GTPase activation instructs adult neural stem cell fate

We next analyzed the effect of optogenetic activation of RhoA and Cdc42 on stem cell lineage commitment. After allowing cells to adhere for 18 hours in proliferation media, cultures were switched to differentiation media and illuminated with blue light at 0.5 μW/mm^2^ for 18 hours, within our previously identified critical mechanosensitive window^16^. Cell fate was analyzed after an additional 6 days, using βIII-tubulin (a.k.a. Tuj1) as a neuronal marker and glial fibrillary acidic protein (GFAP) an astrocytic marker (Fig. 3a). This 18 hour continuous RhoA or Cdc42 activation was sufficient to inhibit neurogenesis and promote astrogenesis in both optoRhoA and optoCdc42 cells (Fig. 3b and c), consistent with our prior reports that stiff matrices or constitutively active Rho GTPases suppress neuronal differentiation^3, 15^. Illumination did not affect wtNSC fate (Supplementary Fig. 2b and c).

**Figure 3.**
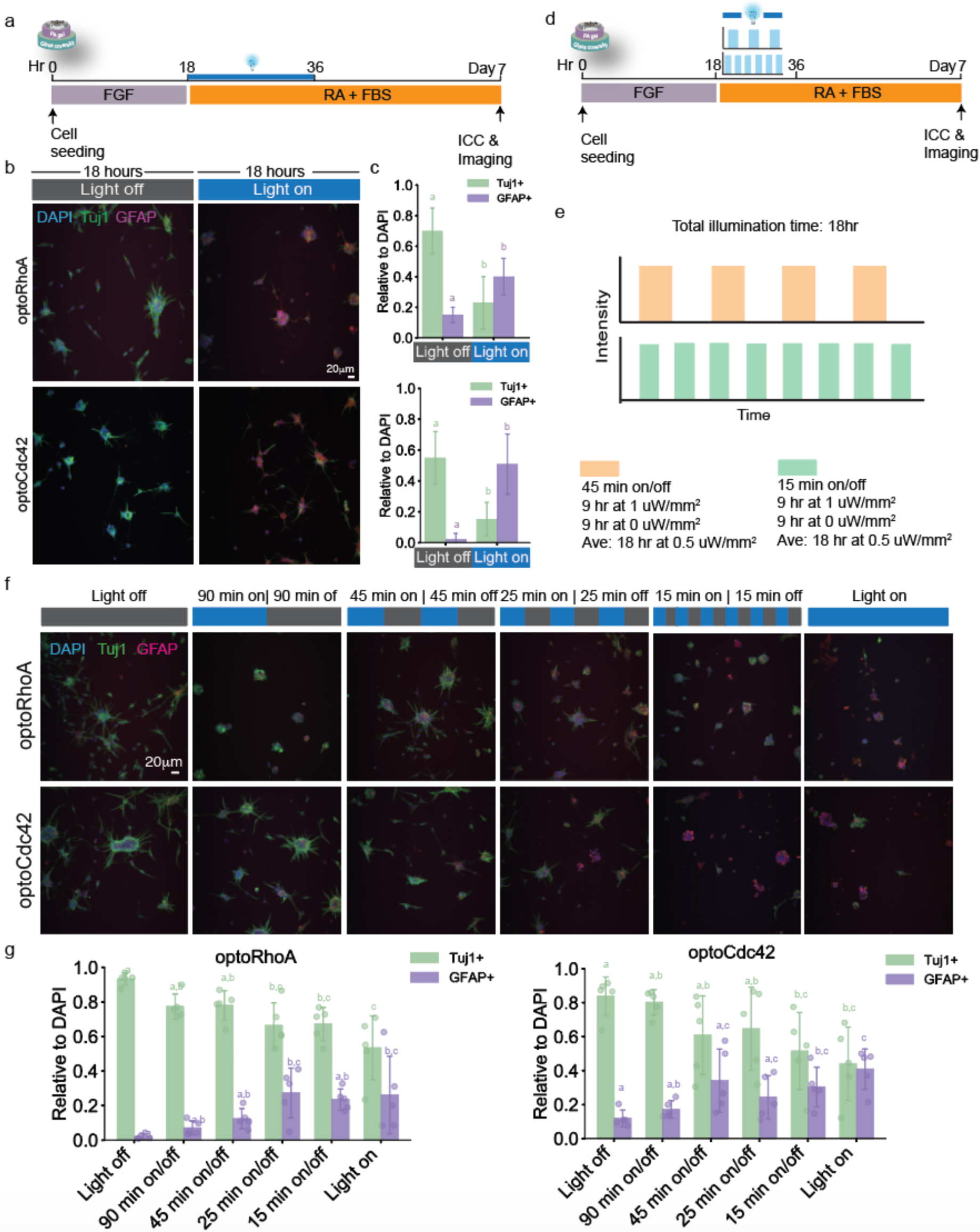
NSC fate is determined by the frequency of activation of Rho GTPases during a critical temporal window. **a**, Media conditions and optogenetic stimulation pattern used to control differentiation of NSCs seeded on hydrogels. **b,** Immunostaining for Tuj1 (green; neurons), GFAP (magenta; astrocytes) or DAPI (blue; nuclei) of optoRhoA or optoCdc42 NSCs after 7 days of differentiation and opto-stimulated for the first 18 hours at a constant light dose of 0.5μW/mm^2^. **c,** Quantification b. Tuj1+/DAPI and GFAP+/DAPI ratios were calculated. Data are represented as mean ± S.D (n=6 biological replicates). **d,** Media conditions and optogenetic stimulation pattern used to control differentiation of NSCs seeded on hydrogels. **e,** Opto-stimulation using LAVA boards allows for independent tunning of frequency and amplitude of the applied light stimuli. **f,** Immunostaining for Tuj1 (green; neurons), GFAP (magenta; astrocytes) or DAPI (blue; nuclei) of optoRhoA or optoCdc42 NSCs after 7 days of differentiation and opto-stimulated for the first 18 hours at different frequencies, keeping an average dose of 0.5μW/mm^2^. **g,** Quantification of f. Tuj1+/DAPI and GFAP+/DAPI ratios were calculated. Data are represented as mean ± S.D (n=6 biological replicates). One-way ANOVA followed by Tukey’s multiple comparison test was used to evaluate statistical significance.

To simulate the Rho GTPase oscillations we observed on stiff and soft gels, we examined the effect of pulsatile RhoA and Cdc42 activation during the critical temporal window (Fig. 3d). We performed a frequency sweep analysis ranging from 15-minute to 90-minute square pulses of light (duty cycle 50%) and doubled the illumination intensity (1 μW/mm^2^) such that light dose was constant for all treatments including the continuous 18 hour light condition (Fig. 3e). While slightly slower than the frequency observed on stiff substrates (Fig. 1f), we chose 15-minute square pulses as the fastest frequency tested considering the clustering/dissociation kinetics of these proteins shown in Figure 2 (∼4 minutes for nearly complete protein clustering and ∼8 minutes for nearly complete dissociation). Strikingly, we found that stem cell fate is sensitive to the frequency of activation of RhoA and Cdc42, with high frequencies inhibiting neurogenesis and promoting astrogenesis to the same extent as continuous illumination (Fig. 3f and g). In contrast, 45-minute or slower square pulses had the same outcome as cells that did not receive light stimulation (Fig. 3f and g). Notably, the proportions of neurons and astrocytes varied continuously with stimulation frequency (Fig. 3g). Overall, these data suggest that Rho GTPases encode their capacity to regulate NSC fate in the frequency of their activation, which in turn is determined by the elasticity of the underlying ECM.

We next investigated whether the dynamic activation of these GTPases was instructing cell fate as opposed to selecting a subpopulation of cells by differentially modulating the proliferation or apoptosis of already committed progenitors. As we previously reported^3, 15^, the majority of individual NSCs have the capacity to generate neurons, astrocytes, and, to a lesser extent, oligodendrocytes. To examine clonal cell behavior at different stimulation frequences, a microisland array technology we previously developed^32^ was used to place a single cell in each island and thereby track single cell fate decisions upon blue light stimulation (Fig. 4a and b). Specifically, using DNA-programmed adhesion, cells were patterned at a density of 1 cell/microisland, illuminated with high (15-minute square pulses) or low (45-minute square pulses) light frequencies over an 18-hour period and analyzed through differential marker expression after 7 days. Wild type NSCs (wtNSCs) gave rise to a distribution of lineage commitments – ∼50% neurons and ∼30% astrocytes) – that did not vary significantly with frequency of illumination (Fig. 4d and e). In contrast, upon illumination of optoRhoA and optoCdc42 cells for 18 hours at different frequencies, the proportion of neurons vs. astrocytes was strongly dependent on stimulation frequency, with microislands harboring 50-70% neurons upon low-frequency stimulation and, strikingly, 95-100% astrocytes upon high-frequency stimulation (Fig. 4c-e). Of note, among microislands that were 100% neuronal or 100% astrocytic, the former had more cells than the astrocytic islands, consistent with previous reports that neuronal but not astrocytic-committed progenitor cells undergo additional proliferation after fate commitment^33^. This explains the even more dramatic inhibition of neurogenesis and promotion of astrogenesis seen in this clonal assay compared to the bulk-population assays (Fig. 3b and f) upon RhoA or Cdc42 activation. This clonal analysis thus reveals high-frequency Rho GTPase activation shifts most stem cells towards astrocytic fates, with a small proportion of neuronal committed cells.

**Figure 4.**
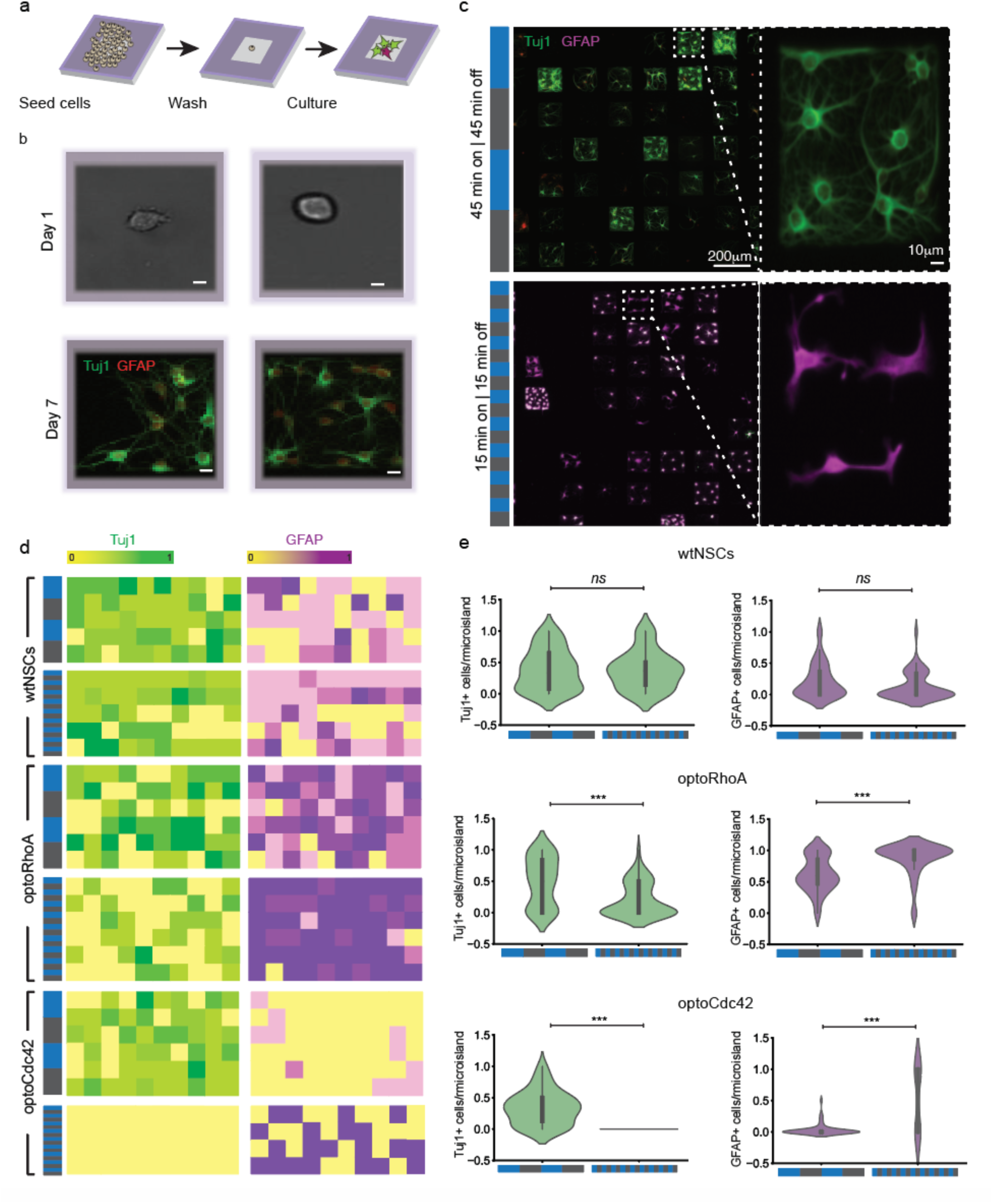
High-frequency Rho GTPases activation promotes astrogenesis through an instructive mechanism. **a,** Schematic representation of NSCs seeding process onto single-cell microisland arrays. **b,** Live-cell images (Day 1) from 2 representative microislands and the corresponding immunofluorescence staining (Day 7) for Tuj1 (green; neurons), GFAP (magenta; astrocytes) or DAPI (blue; nuclei). Scale bar 10 μm. **c,** Immunofluorescence staining for Tuj1 (green; neurons), GFAP (magenta; astrocytes) or DAPI (blue; nuclei) of optoRhoA cells seeded in microislands and differentiated for 7 days. **d,** Graphs show heatmaps indicating the proportion (in a 0-1 scale) of Tuj1+ or GFAP+ cells relative to DAPI in each microisland, for optoRhoA, optoCdc42 or wtNSCs illuminated at different frequencies. **e,** Violin plots showing the distribution of Tuj1+/DAPI and GFAP+/DAPI ratios per microisland. Student’s t-test was used to evaluate statistical significance. \*\*\**P* < 0.005, \*\**P* < 0.01, \**P* < 0.05. ns, not significant.

### Frequency-dependent activation of Rho GTPases controls downstream actin dynamics and mimics the effects of substrate stiffness on cytoskeletal organization

We have previously shown that the downstream contractile and cytoskeletal machinery activated by RhoA and Cdc42 is necessary to mediate stiff substrate inhibition of neurogenesis^15^. To determine whether and how frequency-dependent lineage commitment depends on actin assembly, we seeded optoRhoA and optoCdc42 NSCs on compliant (520 Pa) substrates and illuminated at low (45-minute square pulses) or high frequencies (15-minute square pulses) in the presence of cytochalasin D (Cyto D) to inhibit actin polymerization during the critical 18-hour window (starting 18 hr post-seeding and finishing 36 hr post-seeding) (Supplementary Fig. 3a). This inhibition during the lineage commitment window rescued neurogenesis and suppressed astrogenesis in both optoRhoA and optoCdc42 cells stimulated at high frequencies (Supplementary Fig. 3b and c). We next examined the immediate, short-term effects of RhoA or Cdc42 pulsatile activation on actin dynamics. optoRhoA and optoCdc42 NSCs were engineered to express LifeAct-mVenus to allow for live-cell tracking of actin dynamics^34^ (Supplementary Fig. 3d). A 15-minute pulse of blue light on optoRhoA or optoCdc42 NSCs triggered fast actin disassembly from filopodia-like protrusions and a reorganization around the cortical region and perinuclear compartment (Supplementary Fig. 3e and f), followed by retraction of these protrusions, consistent with increased cellular contractility (Supplementary Fig. 3g). Moreover, clusters of RhoA and Cdc42 formed during these short light pulses and co-localized with actin bundles both in the plasma membrane as well as in the perinuclear region (Supplementary Fig. 3h). Altogether, this data indicates that activation of Rho GTPases triggers fast re-organization of actin bundles within different subcellular regions, both the cortical area and perinuclear compartment, and that actin polymerization is necessary to mediate the effects of oscillatory activation of RhoA and Cdc42 on cell fate (Supplementary Fig. 3i).

To further explore the effects of Rho GTPases pulsatile activation on the temporal dynamics of actin cytoskeleton of NSCs, we analyzed the speed of actin subcellular shuttling via live-cell imaging following the design described above (Supplementary Fig. 3d). We found that optogenetic stimulation of RhoA or Cdc42 for the first 15 minutes decreased shuttling speed compared to non-illuminated cells, suggesting that activation of RhoA or Cdc42 immobilizes a pool of cytoplasmic actin (Supplementary Fig. 4a and b). Furthermore, high-frequency stimulation rendered low actin shuttling velocity, whereas low-frequency stimulation allowed for higher shuttling speed during the off times (Supplementary Fig. 4c). We further compared these effects with the dynamics of actin cytoskeleton on wtNSCs exposed to substrates of different stiffnesses. Interestingly, we found that as early as 4 hours post-seeding there is a trend towards higher actin velocity in cells seeded on soft substrates compared to stiff, which becomes statistically significant at 18 hours post-seeding (Supplementary Fig. 4d and e). These results agree with our previous demonstration that NSCs adapt to the mechanical properties of the substrate, exhibiting greater elastic modulus when cultured on stiff substrates, which suggests higher cytoskeletal tension^15^. Our data indicate that, as with cell fate instruction, high-frequency activation of RhoA or Cdc42 mimics the effects of stiff matrices on actin cytoskeleton dynamics in NSCs, whereas low-frequency activation phenocopies NSCs exposed to soft substrates.

Taking together these findings, we hypothesize that, when exposed to soft substrates, most of the Rho GTPases in these cells will exhibit scarce pulses of activation, leading to the formation of less stable, presumably shorter, actin filaments, rendering a more fluid cytoskeletal network (Supplementary Fig. 4f). On the other hand, higher extracellular tension increases the frequency of activation of Rho GTPases, triggering stable actin filament formation, therefore rendering a more static cytoskeletal organization (Supplementary Fig. 4f). Moreover, actin polymerization is necessary to mediate the effects of frequency-dependent activation of RhoA and Cdc42 on NSC fate commitment, highlighting the critical role of actin dynamics for cell fate instruction.

### Frequency-dependent oscillatory activation of Rho GTPases controls SMAD1/5 phosphorylation and nuclear translocation dynamics in NSCs

We next examined how pulsatile activation of RhoA and Cdc42 and concomitant modulation of actin cytoskeleton dynamics are further transduced downstream into signals that irreversibly determine stem cell commitment. We first investigated effects on β-catenin signaling, as we have mechanistically linked the inhibition of this pathway in NSCs to stiffness-dependent reduction in neurogenesis^16^. We used a β-catenin reporter composed of luciferase placed under the control of a β-catenin responsive promoter (6xTCF) into NSCs^16^; however, we did not detect any significant differences in luciferase activity upon Rho or Cdc42 activation during the critical temporal window, suggesting that a different mechanism may be involved (Supplementary Fig. 5a).

Rho GTPases have been shown to cross-talk with the transforming growth factor beta (TGF-β)/bone morphogenetic protein (BMP) signaling network, which controls the activation of the SMAD family of transcription factors ^35–41^. We and others have shown that these pathways are key regulators of neural stem cell differentiation during both embryogenesis and adulthood^40–43^. In addition, RhoA has been shown to mediate TGFβ-induced smooth muscle differentiation by modulating SMAD activity^37^. We therefore investigated whether the SMAD pathway was playing a role in RhoA/Cdc42-mediated cell fate instruction. We seeded wtNSCs on compliant or rigid substrates and measured SMAD1/5 phosphorylation during the critical mechanosensitive time window. Remarkably, the proportion of cells positive for nuclear pSMAD1/5 was significantly higher on cells exposed to stiff substrates compared to those on soft matrices (Supplementary Fig. 5b and c), indicating that matrices with higher elastic modulus promote activation of the SMAD1/5 pathway.

We next asked whether dynamic Rho GTPase activation within the mechanosensitive temporal window modulates SMAD1/5 activity. optoRhoA, optoCdc42, or wtNSCs were seeded on soft substrates and illuminated either constantly or at different frequencies for a total of 90 minutes, during the critical temporal window. Interestingly, we found that constant light stimulation of RhoA and Cdc42 triggered phosphorylation and nuclear translocation of SMAD1/5 (Fig. 5a-c). In addition, low frequency oscillation likewise induced SMAD1/5 phosphorylation and nuclear translocation (Fig. 5a-c), where 45-minute in the dark followed by a 45-minute illumination induced significant SMAD1/5 phosphorylation and nuclear translocation compared to the dark control (Fig. 5a-c). However, the inverse sequence (a 45-minute light pulse followed by a 45-minute dark pulse) did not yield noticeable SMAD1/5 phosphorylation or nuclear translocation, indicating a 45-minute gap in Rho GTPase activation is sufficient for SMAD1/5 activity to decay to background levels. In contrast, a 90-minute total illumination at a high, 15-minute on/off frequency, led to high levels of SMAD1/5 activity, regardless of illumination phase (Fig. 5a-c). These results indicate that high-frequency Rho GTPase activation leads to persistent, whereas low-frequency activation leads to intermittent, SMAD1/5 phosphorylation and nuclear localization.

**Figure 5.**
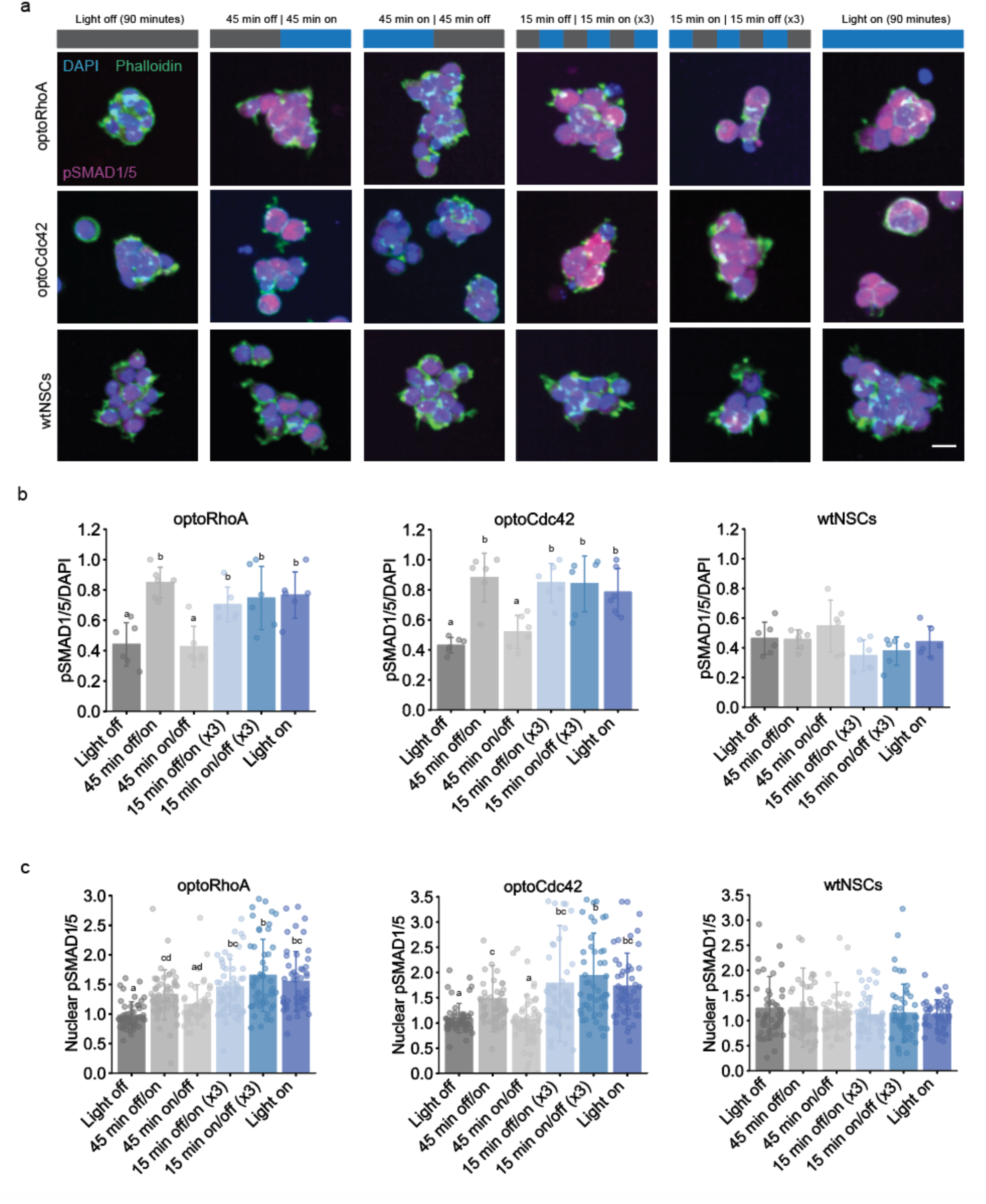
SMAD1/5 phosphorylation and nuclear localization is modulated by the frequency of activation of Rho GTPases. **a,** Immunostaining for pSMAD1/5 Ser463/465 (magenta), phalloidin (green) and DAPI (blue, nuclei) in NSCs seeded on soft PA hydrogels and illuminated at the indicated frequencies for a total of 90 minutes. Average light dose for all conditions was 0.5μW/mm^2^. Scale bar 10 μm. **b,** Quantification of a. pSMAD1/5+/DAPI ratios were calculated for each cell type. Data are represented as mean ± S.D (n=6 biological replicates). One-way ANOVA followed by Tukey’s multiple comparison test was used to evaluate statistical significance. **c,** Quantification of a. Using Fiji^60^, masks of cell nuclei were generated to segment nuclear from cytoplasmic signal. Mean pSMAD1/5 intensity per cell was calculated in the segment cytoplasmic and nuclear regions. Nuclear/cytoplasmic ratios are plotted for each cell type. Data are represented as mean ± S.D (n=50 cells per condition). One-way ANOVA followed by Tukey’s multiple comparison test was used to evaluate statistical significance.

It was recently shown in human embryonic stem cells (hESCs) that increased cellular contractility triggers SMAD1/5 phosphorylation, an effect that persisted in the presence of canonical inhibitors of TGF-β and BMP receptors (SB-431542 and LDN-193189), suggesting that non-canonical, receptor-independent mechanisms may be involved in this mechanotransduction^44^. We thus likewise explored whether the effects of RhoA or Cdc42 activation on SMAD1/5 phosphorylation in NSCs were mediated by TGF-β/BMP receptors. We treated the opto-stimulated cells with dual SMAD inhibitors (SB-431542 and LDN-193189), and found that these inhibitors did not block SMAD1/5 phosphorylation (Supplementary Fig. 5d and e), suggesting that Rho GTPase activation may trigger SMAD1/5 phosphorylation via a mechanism independent of TGF-β/BMP receptors.

We next investigated whether Rho GTPase-triggered actin cytoskeleton reorganization was necessary for SMAD1/5 phosphorylation. Blocking actin cytoskeleton polymerization with Cyto D inhibited the phosphorylation of nuclear SMAD1/5 in response to RhoA or Cdc42 activity (Fig. 6a and b), indicating that an intact actin cytoskeleton is necessary for SMAD1/5 phosphorylation downstream of dynamic Rho GTPase activation.

**Figure 6.**
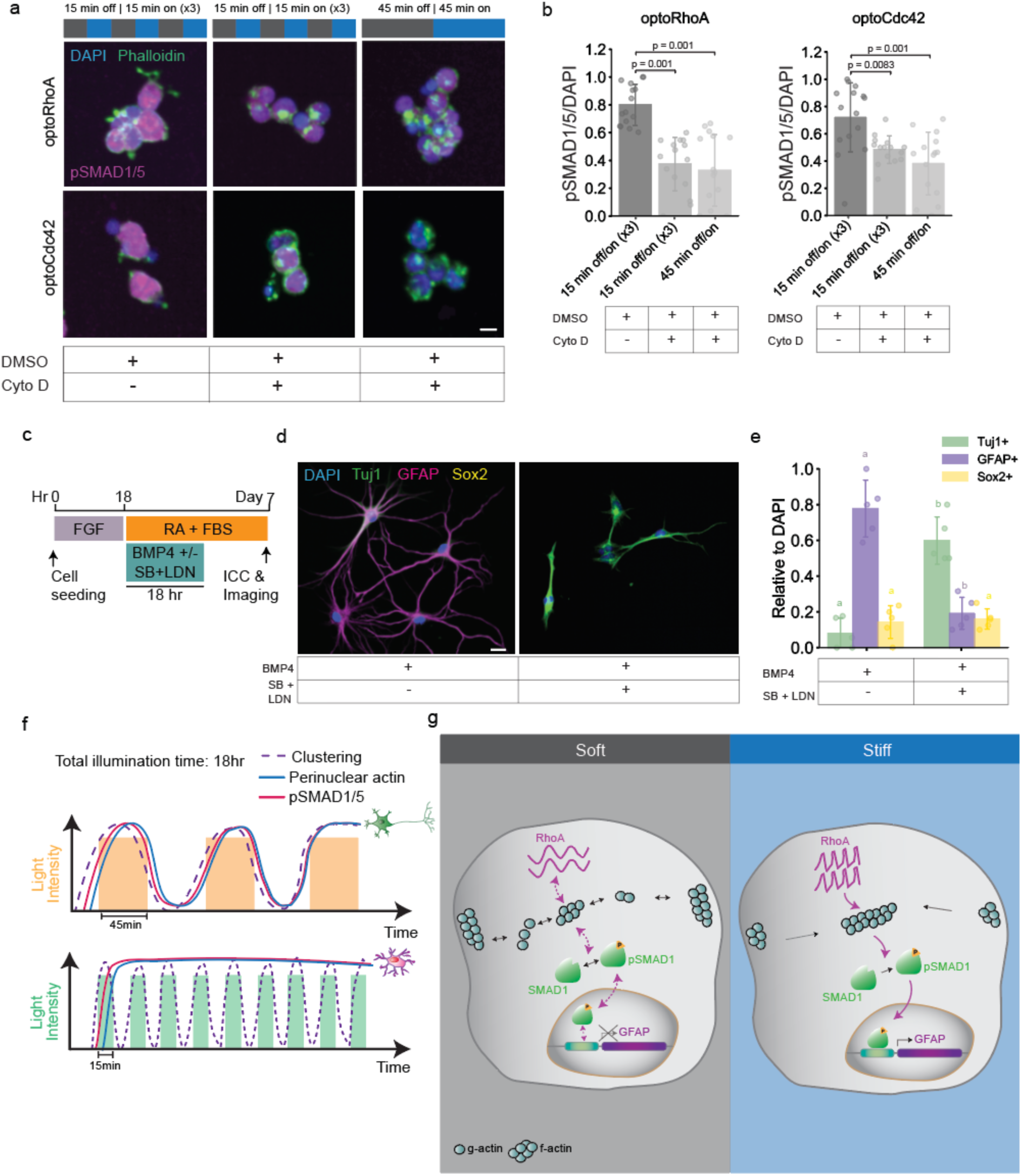
Actin-dependent activation of SMAD1 signaling pathway mediates the effects of dynamic activation of RhoA and Cdc42 on cell fate. **a,** Immunostaining for pSMAD1/5 Ser463/465 (magenta), phalloidin (green) and DAPI (blue, nuclei) in NSCs seeded on soft PA hydrogels and illuminated at the indicated frequencies for a total of 90 minutes in the presence of Cyto D or vehicle control (DMSO). Average light dose for all conditions was 0.5μW/mm^2^. Scale bar 10 μm. **b,** Quantification of a. pSMAD1/5+/DAPI ratios were calculated for each cell type. Data are represented as mean ± S.D (n=10-15 biological replicates). One-way ANOVA followed by Tukey’s multiple comparison test was used to evaluate statistical significance. **c,** Media conditions and optogenetic stimulation pattern used to control differentiation of NSCs. Modulators of canonical SMAD pathway were used as indicated. Pathway was activated with recombinant human BMP-4 (100 ng/ml, R&D Systems), and/or inhibited with LDN-193189 (‘LDN’, 100 nM, R&D Systems) or SB-431542 (‘SB’, 10 μm, R&D Systems). **d,** Immunostaining for Tuj1 (green, neurons), GFAP (magenta, astrocytes), Sox2 (neural progenitors, yellow) and DAPI (blue, nuclei) in NSCs treated as indicated for a total of 18 hours and fixed after 7 days of differentiation. Scale bar 10 μm. **e,** Quantification of e. Tuj1+/DAPI, GFAP+/DAPI and Sox2/DAPI ratios were calculated. Data are represented as mean ± S.D (n=5 biological replicates). Student’s t-test was used to evaluate statistical significance. **f,** Frequency-dependent activation of Rho GTPases renders differential effects on cytoskeletal dynamics and SMAD1/5 activation in NSCs. **g,** Proposed model for how activation of Rho GTPases at a high frequency renders NSC fate commitment into astrocytes by persistent activation of the SMAD1/5 pathway.

### Actin-dependent SMAD1 activity mediates stem cell fate decisions driven by pulsatile Rho-GTPases activation and substrate stiffness

We next examined whether SMAD1/5 is necessary to mediate the effect of dynamic Rho GTPase activation on fate commitment. Western Blot and immunocytochemistry analyses readily detected SMAD1 expression in NSCs, whereas SMAD5 could not be detected to the same extent, at least with the antibody used for these assays, suggesting a higher expression of SMAD1 over SMAD5 in these cells (Supplementary Fig. 6a and b). We thus disrupted SMAD1 expression via a shRNA-mediated knockdown (KD) in optoRhoA NSCs. The most effective shRNA tested (“shRNA SMAD1.2”) resulted in a nearly 80% reduction in SMAD1 protein levels (Supplementary Fig. 6c), and we generated stable optoRhoA NSCs SMAD1 KD cells (‘shSMAD1.2-optoRhoA” NSCs). We next subjected these cells, alongside a RNAi control cell line (“shLacZ-optoRhoA” NSCs), to either high- or low-frequency RhoA activation during the critical 18-hour window, then determined their cell fate after 7 days (Supplementary Fig. 6d). As anticipated, control cells (shLacZ-optoRhoA) exposed to high-frequency activation of RhoA showed a marked inhibition of neurogenesis and promotion of astrogenesis compared to cells subjected to low-frequency stimulation of RhoA (Supplementary Fig. 6e and f). However, SMAD1 knockdown eliminated the differences in the proportions of neurons and astrocytes generated between high-frequency activation conditions and low-frequency conditions (Supplementary Fig. 6e and f), indicating that this transcription factor mediates the frequency-dependent RhoA impact on cell fate decisions.

To further elucidate the role of the SMAD1 pathway on fate commitment in wtNSCs, we used BMP4, a canonical activator of this pathway that binds to type I receptors ALK1/2 and ALK3 and type II receptor BMPR2, triggering downstream phosphorylation of SMAD1/5^45, 46^. Recombinant BMP4 treatment of wtNSCs just during the critical 18-hour window strongly shifted cell fate towards astrocytes (Fig. 6c-e). We further found that addition of TGF-β /BMP receptors inhibitors (LDN-193189 and SB431542) reversed these effects of BMP4 treatment (Fig. 6c-e).

Altogether, these results suggest that stiff matrices increase RhoA and Cdc42 activation frequency, which in turn enhances actin assembly needed to trigger persistent SMAD1/5 activation, rendering a shift in cell commitment towards astrocytic fates. In contrast, softer matrices reduce the frequency of RhoA and Cdc42 activation, producing a more dynamic actin and transiently assembled actin cytoskeleton and reducing SMAD1/5 phosphorylation, and neurogenesis. Critically, we found that the effect of matrix stiffness can be phenocopied by optogenetic stimulation of RhoA and Cdc42 at the appropriate frequency (Fig. 6f and g).

## Discussion

Biophysical cues from the extracellular space are actively sensed by cells through dynamic generation of protrusions coordinated by Rho GTPases. These mechanotransduction proteins are molecular switches that cycle between active and inactive forms in response to mechanical inputs in all mechanosensitive cell types, modulating proliferation, protrusion extension and cell migration^47^. Moreover, work from our and other groups has shown that steady activation of Rho GTPases mediate stiffness-instructed cell fate decisions in NSCs and MSCs, indicating that these proteins are also critical mediators of mechanosensitive differentiation^15, 22^. While these studies clearly demonstrate that substrate stiffness influences the *magnitude* of Rho GTPase activation, it is unclear whether the *dynamics* are similarly affected, or whether these dynamic fluctuations can be integrated downstream into persistent gene expression changes that irreversibly instruct fate commitment. Here, we used FRET-based Rho biosensors and biochemical assays to reveal that Rho GTPases exhibit oscillatory activation dynamics in NSCs, and that the frequency of these oscillations is modulated by the elastic properties of the underlying matrix. We hypothesize a working model in which NSCs respond to substrate stiffness by encoding these mechanical signals into the frequency of activation of the Rho GTPases RhoA and Cdc42 (Fig. 6f and g). Substrate elasticity determines the frequency of GTPase activation, which would function as a strategy for sampling ECM mechanics with high spatio-temporal resolution, through modulating the morphology and dynamics of cellular protrusions. In addition, the frequency of oscillatory RhoA or Cdc42 activation during a critical temporal window of mechanosensitivity dictates stem cell fate through a mechanism that involves modulation of actin cytoskeleton dynamics and downstream dynamic activation of SMAD1/5, which promotes an astrocytic over neuronal fate.

Using RhoA biosensors, we revealed that active RhoA fluctuates at a low frequency in NSCs seeded on soft substrates, with peaks repeating every ∼45-60 minutes, and this frequency is approximately four times higher when cells are seeded on stiff matrices (∼5-10 minute periods). Spontaneous cyclic Rho GTPase activation has been predicted through diverse mathematical models, and has been experimentally observed in fibroblasts^48^, osteosarcoma cells^4^ and xenopus embryos^49^. Osteosarcoma cells in particular have been recently found to exhibit spontaneous pulsatile activation of Rho GTPases, with a period in the order of ∼5 minutes, which agrees in order of magnitude with our data shown here for NSCs seeded on stiff substrates^4, 12^. Oscillatory protein activation dynamics have also been recently described for other members of the Ras family of GTPases in breast cancer cells, in which stochastic Ras-associated protrusions are spontaneously generated, acting as a pacemaker for triggering downstream pulses of ERK activation^10, 50^.

While a number of stem cells are known to undergo mechanosensitive lineage commitment via mechanisms involving Rho GTPases^22, 51, 52^, we provide to our knowledge the first demonstration that Rho undergoes stiffness-dependent dynamic fluctuations and that these fluctuations are important for cell fate determination. Given that Ras and Rho GTPase oscillations have been recently described for other cell types, it is tempting to speculate that frequency-modulated oscillations in Rho GTPase activity in response to substrate stiffness described here for NSCs could represent a widespread mechanism shared among other mechanosensitive proteins in diverse cellular contexts. As has been suggested before^12^, a functional advantage of oscillatory activation dynamics of these GTPases is that they offer a mechanism for cells to continuously probe their environment via dynamic cell contractions and mechanosensitive focal adhesions, and therefore more efficiently adapt to dynamically changing extracellular mechanical forces, in this case during the mechanosensitive time window. In addition, localized RhoA activation and downstream contraction pulses, as opposed to whole-cell activation of Rho GTPases, enable local sensing of dynamic extracellular cues while preventing global contraction of the entire cell, which could otherwise lead to detachment from its matrix.

We revealed that tuning the frequency of pulsatile optogenetic activation of RhoA and Cdc42 within the first few hours of differentiation induces the same differentiation profile obtained by varying substrate stiffness, with low-frequencies resembling soft substrates and high-frequencies phenocopying the effects of stiff matrices. This frequency-dependent oscillatory activation of Rho GTPases modulates actin cytoskeleton dynamics and determines downstream SMAD1/5 phosphorylation and nuclear translocation pattern. SMADs are a family of transcription factors that lie at the core of the transforming growth factor beta (TGFβ) and bone morphogenetic protein (BMP) pathways, with critical roles in embryogenesis such as for patterning of neuroectoderm during cerebral cortex development^53, 54^, as well as in later stages of embryogenesis in which they have been implicated as promoters of astrocytic differentiation from radial glial progenitor cells^41^. In the adult brain, we and others have shown that the BMP-SMAD pathway inhibits neurogenesis^42, 43, 55^, and it has been recently linked to astrocytic promotion, though the underlying mechanism remains to be elucidated^39, 40^. Our results support a role of SMAD1/5 downstream of RhoA and Cdc42 activation during a critical window of mechanosensitivity, in instructing astrogenesis over neurogenesis in NSCs.

Modulation of temporal dynamics in SMAD1/5 phosphorylation and nuclear translocation in response to the frequency of pulsatile optogenetic activation of Rho GTPases suggests that a persistent nuclear localization of pSMAD1/5 is required to shift cell commitment from neuronal to astrocytic fates, whereas sparse pulses of SMAD1/5 activation were not sufficient to instruct cell fate. Temporal modulation of SMAD activation has also been described for fibroblasts, wherein serum treatment triggers sparse oscillations of SMAD activation (∼2 hr period)^56^. Foundational research investigating the temporal regulation of transcription factor dynamics demonstrated that activation of several factors including NF-kappaB in lymphocytes is sensitive to the frequency of intracellular calcium concentration oscillations, driving phenotypic outcomes ranging from interleukin secretion to differentiation^57, 58^. Recent work also showed that sustained versus pulsatile activation of NF-kappaB in macrophages differentially affected target gene expression^59^.

Overall, our data strongly support a role for the Rho GTPase activation dynamics in processing stiffness cues and instructing mechanosensitive stem cell differentiation. These effects can be phenocopied by optogenetic stimulation of RhoA and Cdc42 at the appropriate frequency. These findings advance our fundamental understanding of how ECM mechanics influence cell fate and could further aid in the development of novel strategies to control the timing of biophysical differentiation cues and modulation of Rho and SMAD signaling pathways for improved stem cell lineage manipulation, which remains an essential challenge for regenerative therapies.

## Acknowledgments

We would like to thank Dr. Mary West, Deepa Sridharan, Holly Aaron and Feather Ives for their microscopy advice and support. This work was supported by the NIH (R01NS074831 to S.K. and D.V.S.) and a Siebel Postdoctoral Fellowship to R.G.S. Imaging conducted at the CRL Molecular Imaging Center, RRID:SCR_017852, was supported by the Helen Wills Neuroscience Institute.

## Author contributions

R.G.S. designed and performed the experiments, analyzed, plotted, and interpreted the data, and wrote the manuscript. M.S. performed experiments and contributed to data interpretation. S.K. and D.V.S. supervised the project, providing insights in experimental design and data interpretation, revised and edited the manuscript.

---

**Supplementary Figure 1.**
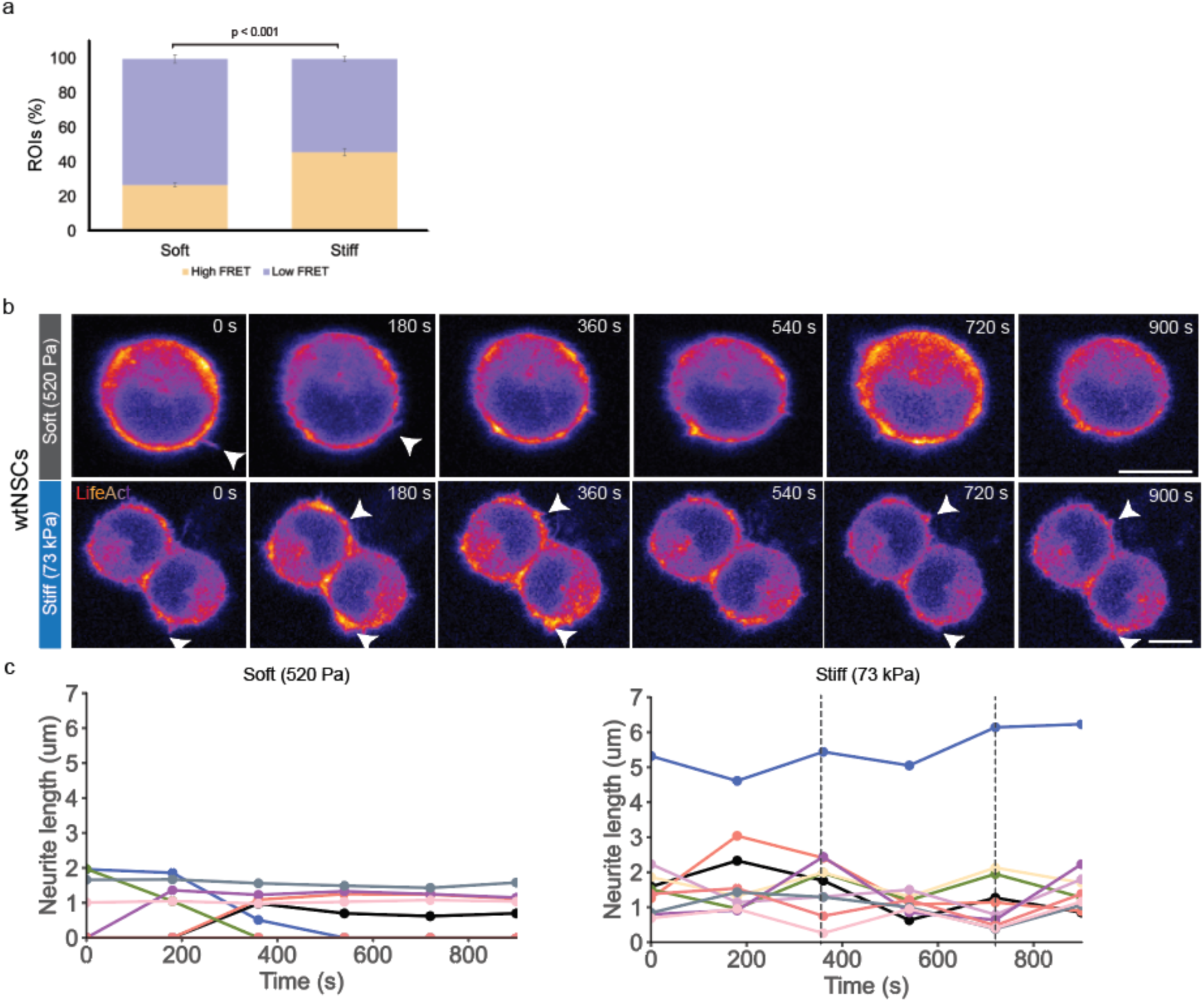
Stiffness-dependent frequency-modulated oscillatory RhoA activation and neurite extension. **a**, Frequency of ROIs per cell with high RhoA activity (average normalized FRET index amplitude ≥ 0.4) or low (average normalized FRET index amplitude ≤ 0.2) was calculated for wtNSCs seeded on soft or stiff substrates. Student’s t-test was used to evaluate statistical significance. **b,** Time-lapse imaging of LifeAct on wtNSCs allows for tracking neurite extension/retraction dynamics on cells seeded on hydrogels of different stiffnesses. Scale bar 10 μm. **c,** Quantification of b. Single-neurite lengths were calculated over time using the simple neurite tracer plugin in Fiji^60^ (n=10 cells per condition).

**Supplementary Figure 2.**
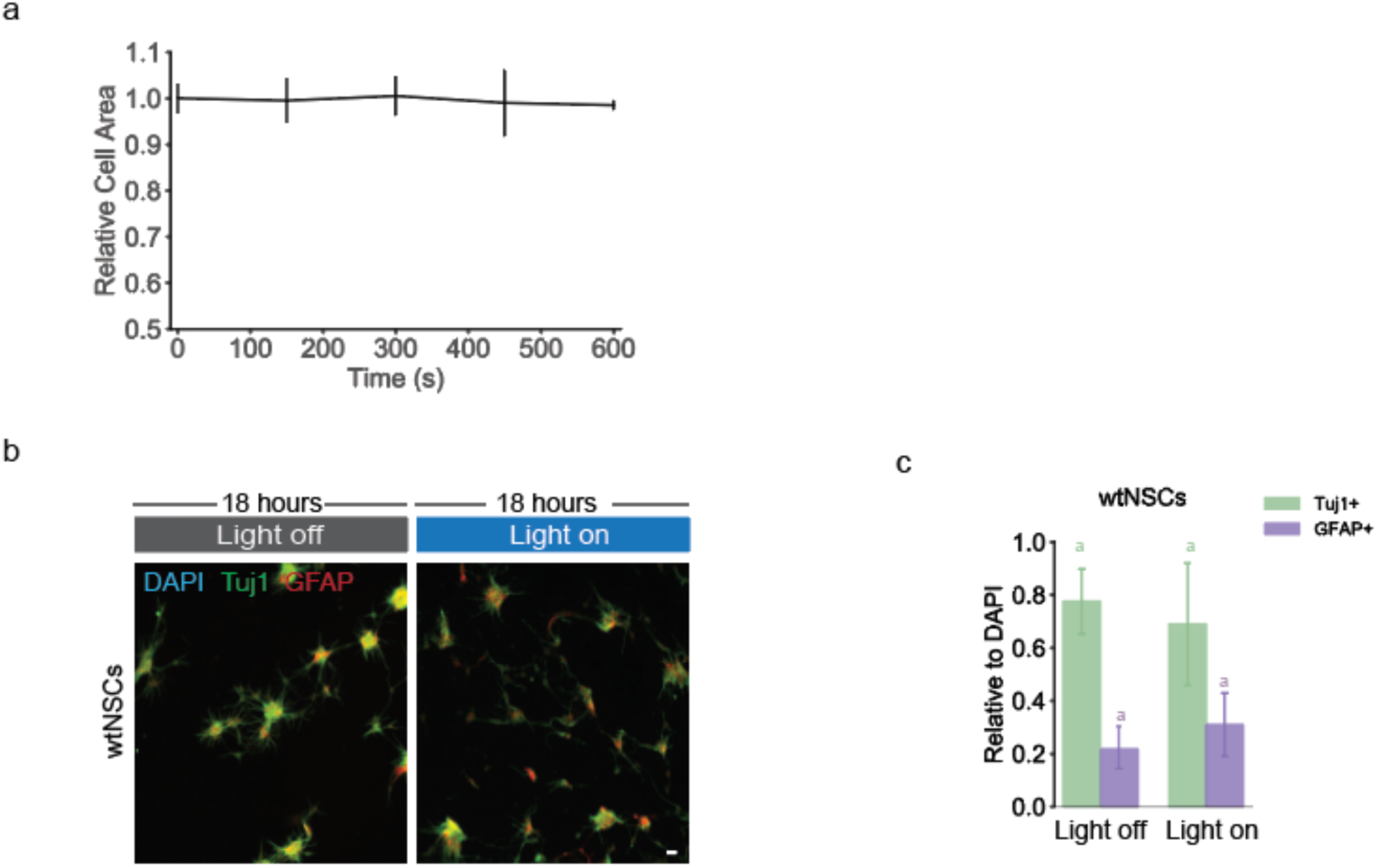
Blue-light stimulation does not affect cell morphology or lineage commitment in wtNSCs. **a,** Contraction of wtNSCs upon light stimulation was determined through live-cell imaging by measuring the cell area at each timepoint with the software Fiji^60^. Resulting cell areas at each timepoint were plotted relative to *t*=0. **b,** Immunofluorescence staining for Tuj1 (green; neurons), GFAP (magenta; astrocytes) or DAPI (blue; nuclei) of wtNSCs cells seeded on soft hydrogels, treated with blue light for 18 hours and fixed after 7 days of differentiation. Scale bar 10 μm. **c,** Quantification of b. Tuj1+/DAPI and GFAP+/DAPI ratios were calculated. Data are represented as mean ± S.D (n=4 biological replicates). Student’s t-test was used to evaluate statistical significance.

**Supplementary Figure 3.**
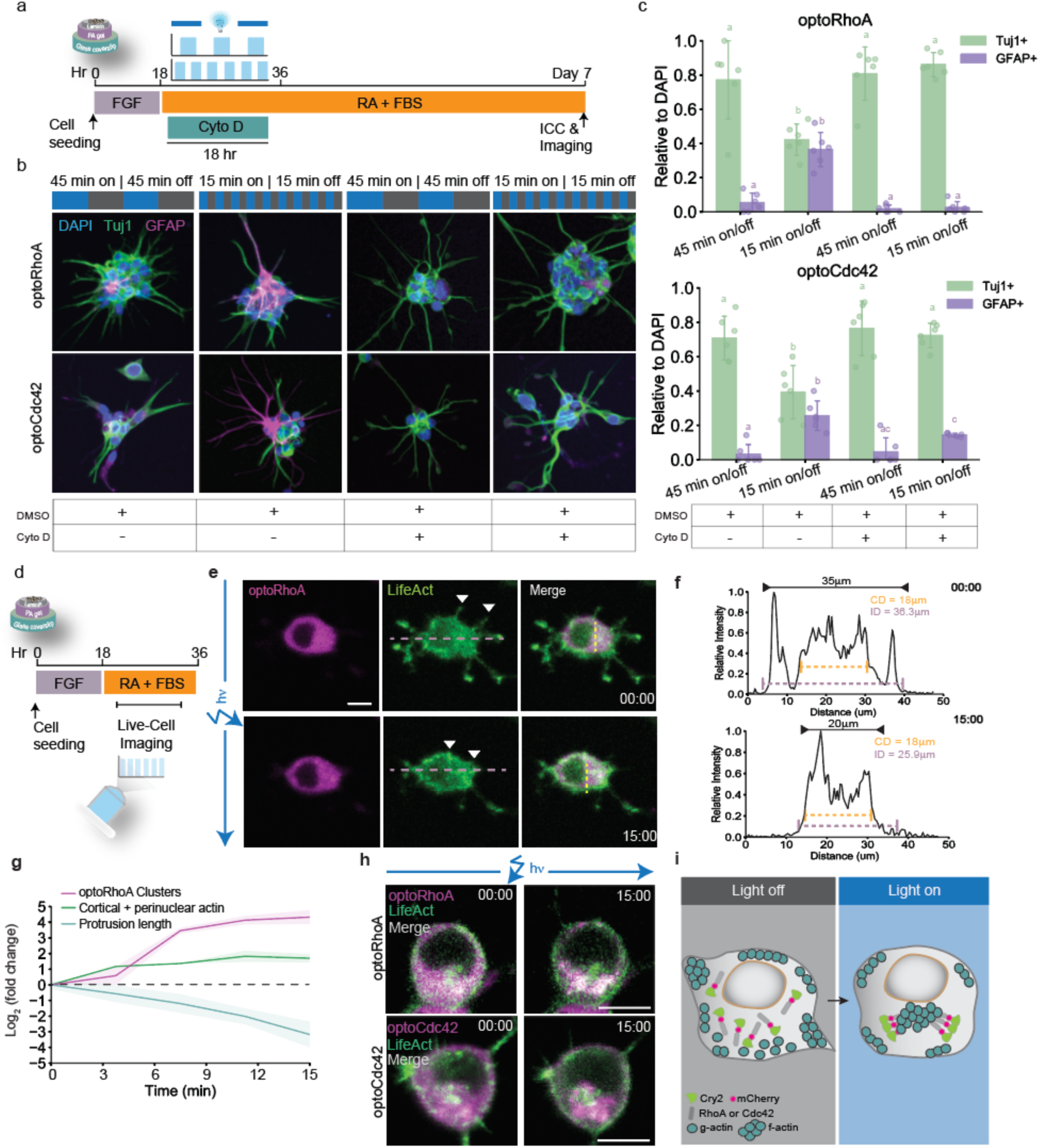
Actin reorganization upon dynamic Rho GTPases activation is necessary to mediate the effects on cell fate decision. **a,** Media conditions and optogenetic stimulation pattern used to control differentiation of NSCs seeded on hydrogels. Where indicated, actin assembly was inhibited with cytochalasin D (Cyto D, 1 μM, Sigma-Aldrich). Control conditions received DMSO vehicle. **b,** Immunofluorescence staining for Tuj1 (green; neurons), GFAP (magenta; astrocytes) or DAPI (blue; nuclei) of optoRhoA or optoCdc42 cells seeded on soft hydrogels, treated with Cyto D or DMSO as indicated and opto-stimulated at different frequencies. **c,** Quantification of b. Tuj1+/DAPI and GFAP+/DAPI ratios were calculated. Data are represented as mean ± S.D (n=6 biological replicates). One-way ANOVA followed by Tukey’s multiple comparison test was used to evaluate statistical significance. **d,** Media conditions and optogenetic stimulation pattern used to control differentiation and perform live-cell analysis of NSCs seeded on hydrogels. **e,** Live-cell tracking of neurite extension dynamics (LifeAct sensor, green) during optogenetic stimulation for 15 minutes of optoRhoA (magenta). Overlay is shown in white. Scale bar 10 μm. **f,** Representative plot of single-cell tracking of cytoplasmic diameter (CD) and inter-neurite distance (ID) measured at *t*=0 and *t*=15 minutes. Morphological measurements were performed with Fiji^60^. **g,** Fold change in number of optoRhoA clusters, actin intensity (cortical + perinuclear) and protrusion length per cell were plotted. Data are represented as mean ± S.D (n=3 biological replicates). **h,** Live-cell tracking of actin dynamics (LifeAct sensor, green) during optogenetic stimulation for 15 minutes of optoRhoA or optoCdc42 (magenta). Overlays are shown in white. Scale bar 10 μm. **i,** Cartoon showing proposed model for actin fiber subcellular reorganization upon optoRhoA or optoCdc42 clustering.

**Supplementary Figure 4.**
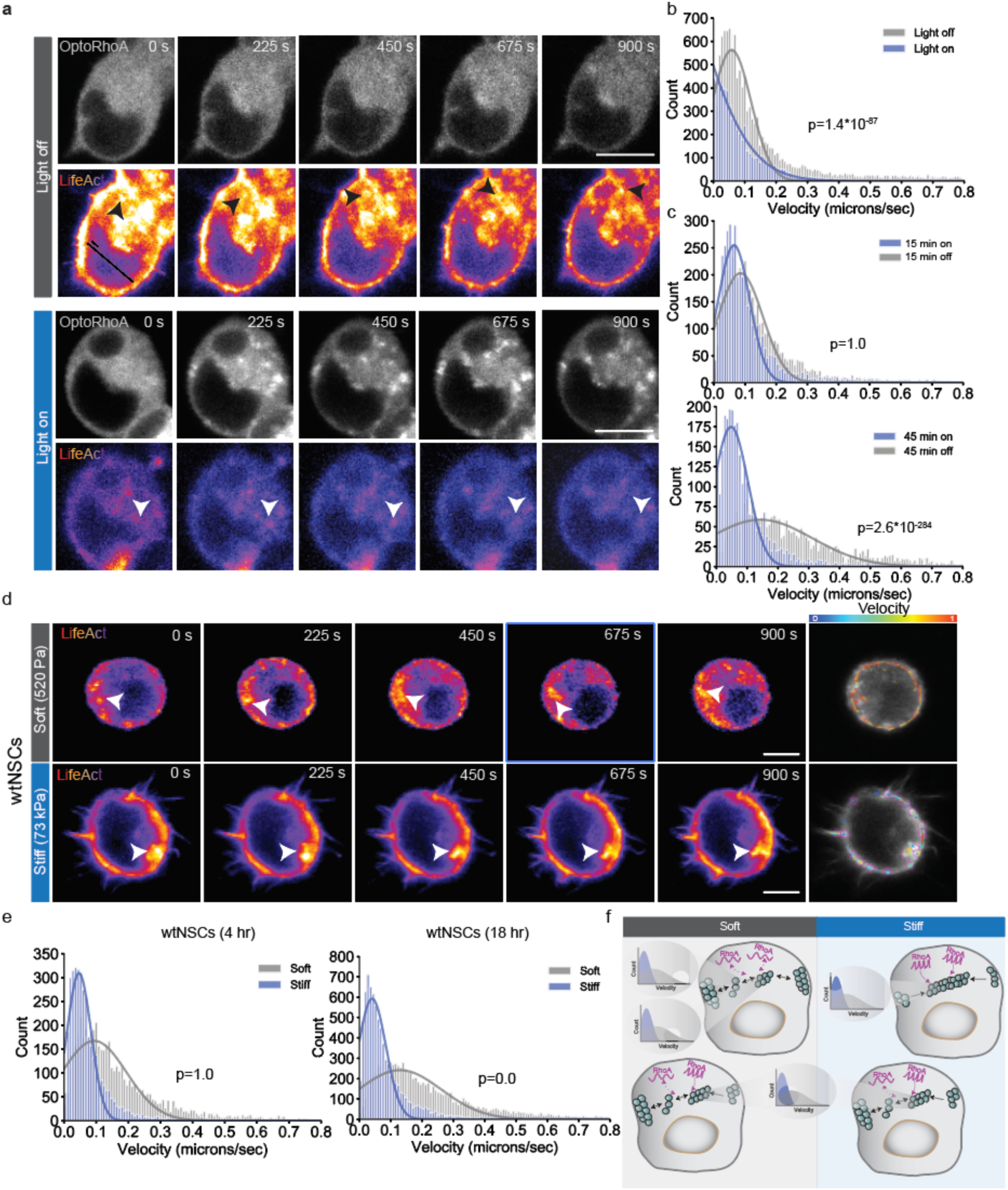
Actin subcellular shuttling dynamics is modulated by the frequency of activation of Rho GTPases. **a,** Time-lapse imaging of LifeAct on optoRhoA to evaluate actin subcellular shuttling dynamics upon opto-stimulation at different frequencies. Scale bar 10 μm. **b,c,** Quantification of a. Single-particle analysis was used to track subcellular dynamics of actin bundles (clusters of ∼500 nm in diameter) using the TrackMate plugin from Fiji^60^. Velocity data for individual particles during the ‘on’ or ‘off’ phases is represented in histograms. Kolmogorov-Smirnov test was used to determine statistical significance **d,** Time-lapse imaging of LifeAct on wtNSCs to evaluate actin subcellular shuttling dynamics on cells seeded on hydrogels of different stiffnesses at 4 and 18-hr post-seeding. Scale bar 10 μm. **e,** Quantification of d. Single-particle analysis was used to track subcellular dynamics of actin bundles (clusters of ∼500 nm in diameter) using the TrackMate plugin from Fiji^60^. Velocity data for individual particles during the ‘on’ or ‘off’ phases is represented in histograms. Kolmogorov-Smirnov test was used to determine statistical significance. **f,** Cartoon illustrating a proposed model for how mechanosensitive, frequency-modulated, oscillatory activation of RhoA can differentially regulate actin cytoskeleton remodeling, leading to differences in subcellular shuttling dynamics.

**Supplementary Figure 5.**
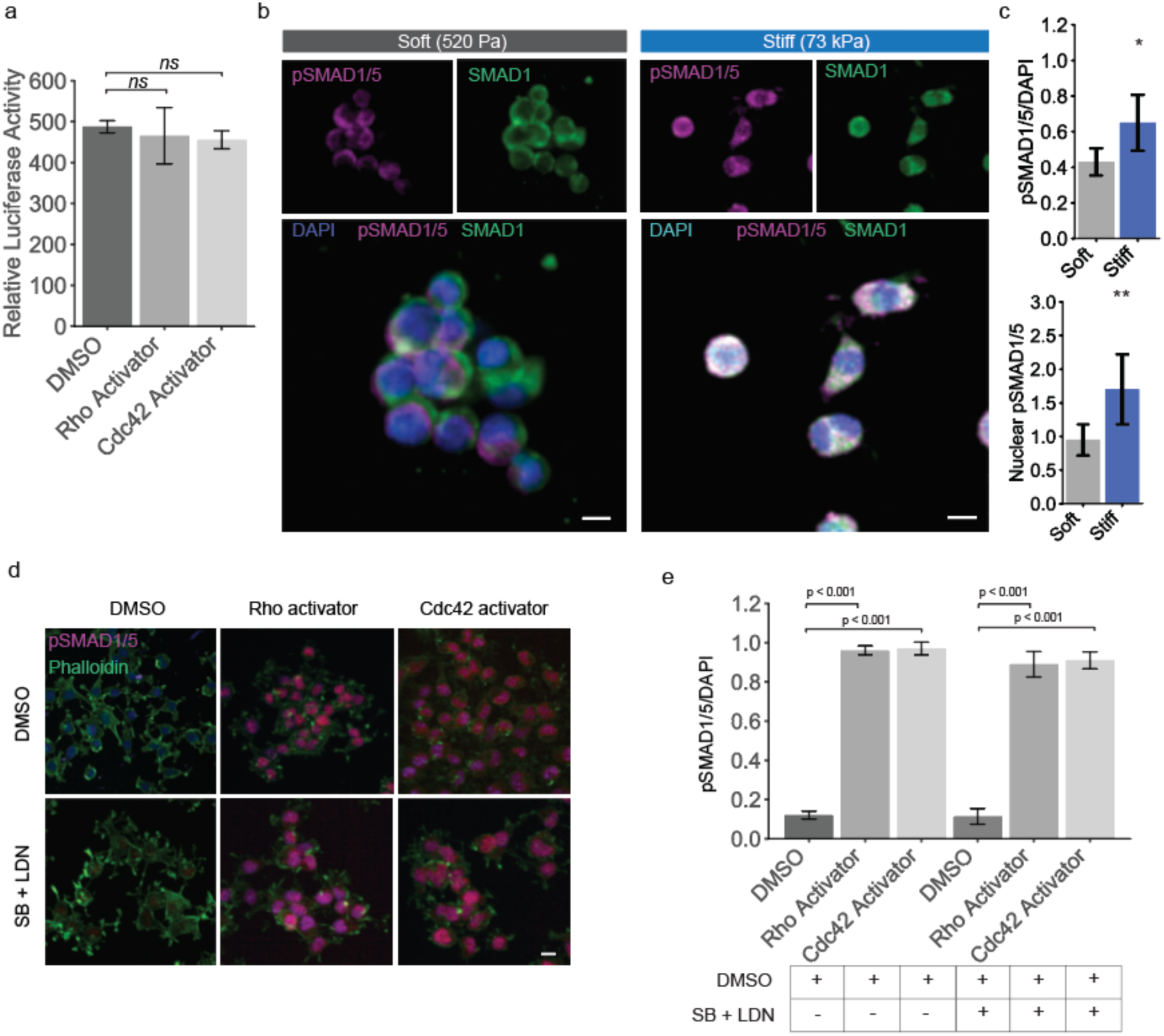
Mechanosensitive activation of SMAD1/5. **a,** Luciferase assay performed on wtNSCs treated as indicated for 18 hours: Rho activator (0.075 units/ml; Cytoskeleton, Inc.), Cdc42 activator (0.75 units/ml; Cytoskeleton, Inc.). One-way ANOVA followed by Tukey’s multiple comparison test was used to evaluate statistical significance. ns, not significant. **b,** Immunostaining for pSMAD1/5 Ser463/465 (magenta), SMAD1 (green) and DAPI (blue, nuclei) in wtNSCs seeded on PA hydrogels of different stiffnesses. Scale bar 10 μm. **c,** (top) Quantification of b. pSMAD1/5+/DAPI ratios were calculated for each cell condition. Data are represented as mean ± S.D (n=6 biological replicates). One-way ANOVA followed by Tukey’s multiple comparison test was used to evaluate statistical significance. **c,** (bottom) Quantification of b. Using Fiji^60^, masks of cell nuclei were generated to segment nuclear from cytoplasmic signal. Mean pSMAD1/5 intensity per cell was calculated in the segment cytoplasmic and nuclear regions. Nuclear/cytoplasmic ratios are plotted for each condition. Data are represented as mean ± S.D (n=50 cells per condition). One-way ANOVA followed by Tukey’s multiple comparison test was used to evaluate statistical significance. \*\*\**P* < 0.005, \*\**P* < 0.01, \**P* < 0.05. ns, not significant. **d,** Immunostaining for pSMAD1/5 Ser463/465 (magenta), Phalloidin (green) and DAPI (blue, nuclei) in wtNSCs treated as indicated for 15 minutes: Rho activator (0.075 units/ml; Cytoskeleton, Inc.), Cdc42 activator (0.75 units/ml; Cytoskeleton, Inc.), LDN-193189 (‘LDN’, 100 nM, R&D Systems) or SB-431542 (‘SB’, 10 μm, R&D Systems). Scale bar 10 μm. **e,** Quantification of d. pSMAD1/5+/DAPI ratios were calculated for each cell type. Data are represented as mean ± S.D (n=6 biological replicates). One-way ANOVA followed by Tukey’s multiple comparison test was used to evaluate statistical significance.

**Supplementary Figure 6.**
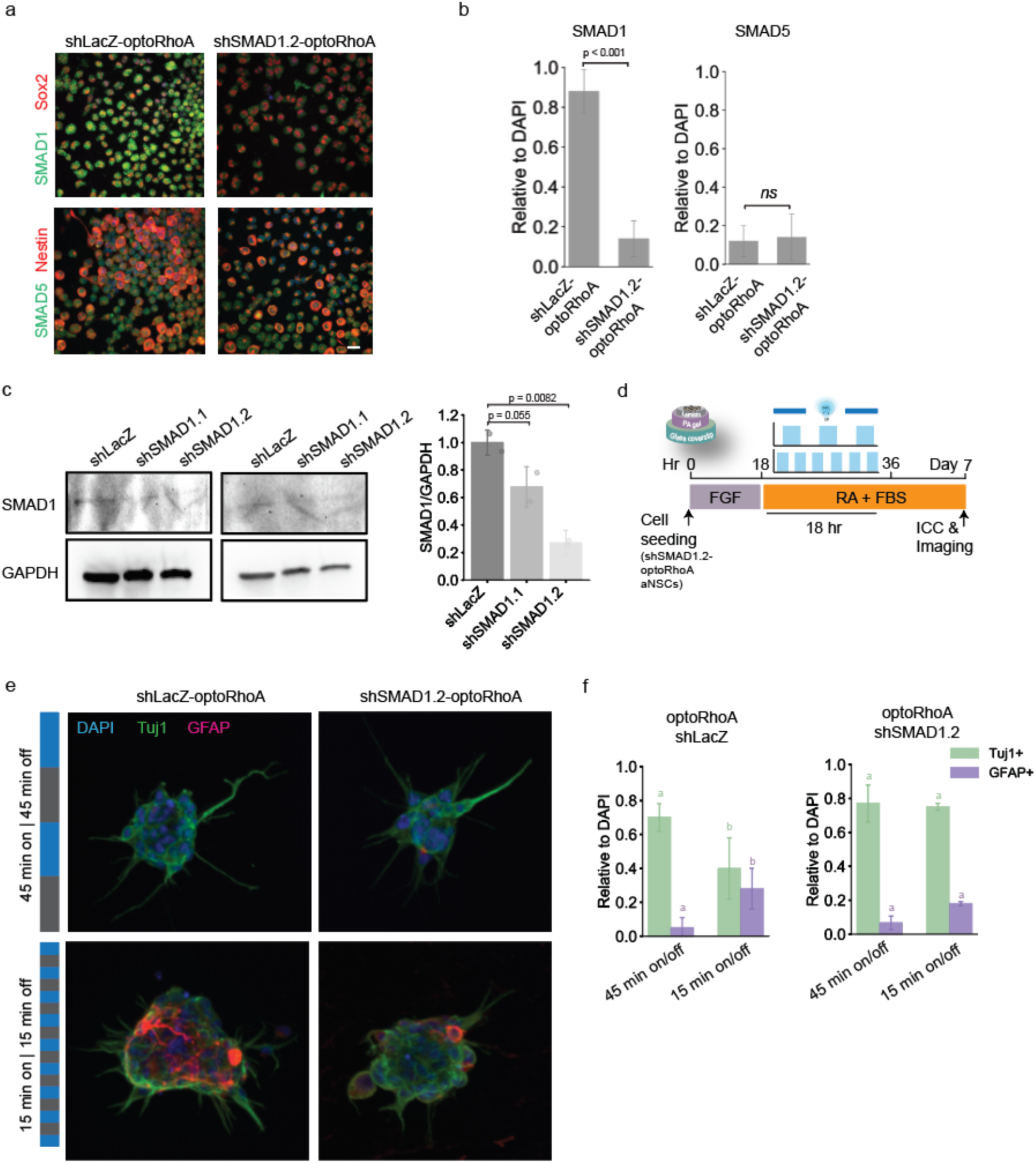
SMAD1 activity mediates fate decisions instructed by Rho GTPases dynamics. **a,** Immunostaining for Sox2 or Nestin (Red), SMAD1 or SMAD5 (green) and DAPI (blue, nuclei) in shSMAD1.2-RhoA or LacZ control to verify reduction in SMAD1 expression through RNAi, and endogenously low SMAD5 expression. Scale bar 10 μm. **b,** Quantification of f. SMAD1+/DAPI and SMAD5+/DAPI ratios were calculated. Data are represented as mean ± S.D (n=2 biological replicates). Student’s t-test was used to evaluate statistical significance. ns, not significant. **c,** Left: Western blot analysis of SMAD1 expression upon RNAi-mediated knock-down 2 different shRNAs targeting SMAD1 (shSMAD1.1 and shSMAD1.2) or LacZ control. Right: Quantification. SMAD1/GAPDH band intensity ratios are shown for each condition. One-way ANOVA followed by Tukey’s multiple comparison test was used to evaluate statistical significance. **d,** Media conditions and optogenetic stimulation pattern used to control differentiation of shSMAD1.2-optoRhoA NSCs seeded on hydrogels. **e,** Immunostaining for Tuj1 (green, neurons), GFAP (magenta, astrocytes) and DAPI (blue, nuclei) in optoRhoA-shSMAD1.2 NSCs seeded on soft PA hydrogels, illuminated at the indicated frequencies for a total of 18 hours and fixed after 7 days of differentiation. Scale bar 10 μm. **f,** Quantification of e. Tuj1+/DAPI and GFAP+/DAPI ratios were calculated. Data are represented as mean ± S.D (n=5 biological replicates). Student’s t-test was used to evaluate statistical significance.

**Supplementary Figure 7.**
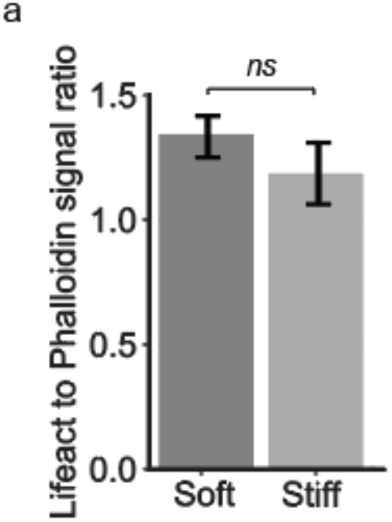
LifeAct-mVenus primarily binds to F-actin in NSCs. **a,** Cells expressing LifeAct-mVenus were fixed and stained for mVenus and Phalloidin (which specifically detects F-actin). LifeAct/Phalloidin intensity ratio was calculated using Fiji^60^. ns, not significant.

## Methods

### Cell culture and differentiation

Adult rat neural stem cells were cultured as described previously^3, 16^. Briefly, cells were cultured in Dulbecco’s modified Eagle’s medium/nutrient mixture F-12 (DMEM-F12, Gibco) with N2 Supplement (Life Technologies) and 20ng/mL fibroblast growth factor (FGF-2, Peptrotech) on tissue-culture polystyrene plates coated, sequentially, with poly-ornithine (10 μg/ml, Sigma-Aldrich) and laminin (5 μg/ml, Invitrogen). For differentiation experiments, cells were seeded on hydrogels of different stiffnesses in proliferation medium for 18 hours. Afterwards, medium was switched to mixed differentiation media (DMEM-F12 with N2, 1 μM retinoic acid, 1% fetal bovine serum). The medium was replaced every 2 days during 7 days of differentiation process.

### Polyacrylamide Hydrogels

Polyacrylamide (PA) hydrogels were produced as described previously^15, 16^. Briefly, acrylamide and bisacrylamide precursor solutions were mixed with APS/TEMED (BioRad) and polymerized onto PlusOne Bind-Silane treated glass coverslips with 0.1% ammonium persulfate and tetramethylethylenediamine. The solution was allowed to polymerize for 15 minutes at room temperature. Laminin functionalization was achieved by incubating gels in 50μg/ml sulfo-SANPAH and irradiating with a UV flood lamp for 8 minutes. The gels were then washed once in PBS, and incubated with 20μg/ml laminin for 4 hours at 37°C.

### Rheological Characterization

Rheological characterization of all hydrogels was performed by shear rheology via a Physica MCR 301 rheometer (Anton Paar) with an 8-mm parallel plate geometry. For frequency sweep measurements, frequency was controlled to be between 50 and 1 Hz at a constant strain (γ = 0.5%). Modulus saturation curve with time was obtained under oscillation with constant strain (γ = 0.5%) and frequency (*f* = 1 Hz). The temperature of the gel solution was controlled (*T* = 37°C) with a Peltier element (Anton Paar). The solvent trap around the sample location was filled with water to prevent sample dehydration. The temperature of the sample was controlled by a Peltier element (Anton Paar).

### Live-cell FRET Imaging and Analysis

The optimized RhoA biosensor (RhoA2G) was acquired from Addgene (plasmid #40179). This sensor consists of the RhoA-binding domain (RBD) of the effector rhotekin that specifically binds guanosine triphosphate (GTP)–RhoA, a mTFP1 donor fluorophore (DF), a 60–amino acid linker, the acceptor fluorophore (AF) Venus, and RhoA^24^. Upon GTP loading, the RhoA portion interacts with RBD, leading to a conformational change and an increase in FRET efficiency. This 2nd generation biosensor system developed by the Pertz lab, was optimized to maximize the emission ratio change between the ON and OFF states of RhoA (achieving 147% ratio change).

The acquisition time for the donor and the FRET channels spanned between 20 and 100 ms with a 20× objective lens using a confocal Zeiss LSM 880 NLO AxioExaminer. Donor and FRET images were acquired sequentially using filter wheels with the following excitation and emission settings: donor channel: 458 laser, 480/40 m emission; FRET channel: 458 laser, 535/30 m emission; acceptor channel: 514 laser, 535/30 emission.

Processing of ratiometric data sets was performed with the FRET analyzer plugin from Fiji^60^. Images were sequentially thresholded on each channel, shade- and background-corrected, masked, and registered before ratios were calculated, as described before for single-chain biosensors^24, 61^. Full scripts for batch processing of time-lapse images with FRET analyzer are available on Github: https://github.com/rosampa/FIJI-FRET-analyzer-batch-processing. Ratio images are color-coded so that warm and cold colors represent high and low biosensor activity, respectively. Particle tracking over time was performed on ROIs of ∼400nm diameter. For frequency calculation, all FRET indexes for each ROI were normalized to t=0. For determination of oscillation frequencies, temporal FRET index profiles were smoothed and detrended to filter out monoexponential decays due to photobleaching, and peaks were counted and averaged to calculate oscillation frequencies, as described before^62^.

### Optogenetic Stimulation

Cells were seeded on PA hydrogels on round coverslips (18mm diameter) inserted on 24-well plates (0030741021, Eppendorf, black-walled with 170µm coverglass bottom), and placed onto LAVA illumination devices kept in standard 37°C tissue culture incubators. In brief, user-defined illumination patterns were uploaded to the LAVA device for independent illumination control of each well^28^. Optogenetic stimulation was achieved with blue light emitted by arrays of 470nm LEDs continuously illuminating hESCs with 0.5 *µ*W mm^-2^ light for the duration of the experiment (18 hrs) or with 1 *µ*W mm^-2^ for frequency sweep analysis.

To assess clustering dynamics in live cells, simultaneous blue light exposure for opto-stimulation and mCherry imaging was accomplished by imaging in both 488-nm and 561-nm laser channels, one exposure per 2-5 s, with the 488-nm laser at 0.5% power using a confocal Zeiss LSM 880 NLO AxioExaminer with a 20× objective lens.

### Single-Cell Microisland Array Preparation and Imaging

#### Micropatterning 20-bp amine-terminated oligonucleotides with positive photoresist

Microisland arrays were manufactured as described before^32, 63^. Briefly, traditional photolithography was used to pattern aldehyde glass substrates with positive photoresist and photoresist-coated aldehyde slides were exposed selectively to UV light (365 nm; 260 mJ cm^−2^) with a mask aligner (Karl Suss MJB 3) using a custom mylar mask (FineLine Imaging). Immediately following photolithography, a 5′-amine–modified, 20-bp oligonucleotide solution prepared in 50 mM sodium phosphate buffer (pH 8.5) was dropcast over the photoresist patterns. DNA oligonucleotides were purchased from both Integrated DNA Technologies, Inc. and Eurofins, and resuspended as 2 mM stocks in molecular biology–grade water. Slides were then heated for 1 hour in a 75°C oven to induce the formation of Schiff bonds (C═N) between the terminal amine on the DNA and the aldehyde on the glass surface. To covalently conjugate the DNA strands to the surface aldehyde groups—thereby converting the hydrolysable Schiff base to single C─N bonds—reductive amination was conducted at room temperature for 15 min in 0.25% sodium borohydride (Sigma-Aldrich) in 1× phosphate-buffered saline (PBS). Upon completion, slides were thoroughly rinsed first with acetone and then DI water, followed by drying with dry nitrogen gas.

#### PA patterns for high-throughput single-cell cultures

Upon completion of DNA patterning, a PA grid was fabricated onto the substrates to enable clonal analysis of thousands of single-cell cultures over the course of differentiation. This was achieved by first photopatterning a large-scale array of 141 μm by 141 μm square features (i.e., “microislands”) arranged with a 200-μm pitch using positive photoresist (Shipley 1813), as described before^32, 63^. Subsequently, a 10% PA solution containing TEMED and APS was dropcast immediately over the photoresist features, and a Gel Slick (Lonza)–treated glass coverslip was used to spread out the PA solution over the entire DNA-patterned substrate. Last, the photoresist defining the PA patterns and protecting the DNA were removed by dissolving in acetone. The slide was rinsed with DI water, dried with dry nitrogen gas, and stored under vacuum.

#### Single-NSC patterning and imaging

Before cell-patterning experiments, DNA-patterned substrates were blocked with 2% BSA in PBS for 1 hour to minimize nonspecific cell attachment. Oligo-labeled NSCs were resuspended in 2% BSA at 4 × 10^7^ cells ml^−1^, and 20 μl was injected into the PDMS flow cell. The high cell concentration ensured that the entire DNA-patterned area was covered with oligo-labeled NSCs. Cells were then cycled by pipetting 5 μl of the cell suspension into the inlet of the flow channel and removing 5 μl from the outlet. This action was repeated 10 to 20 times to increase the chance of hybridization between the cell-tethered oligos and complementary, surface-tethered oligos. Unpatterned cells were washed away with PBS. Upon complete cell patterning, 250 μl of the appropriate culture media supplemented with 10 μg ml^−1^ laminin was added to each well, and the slide was cultured for 7 days with half media changes (minus laminin) every other day to prevent cells from lifting off the surface.

For opto-stimulation experiments with cells seeded on single-cell microisland arrays, blue light stimulation was accomplished by imaging through a GFP filter set (472/30nm) using a ImageXpress Micro System (Molecular Devices) with environmental control (37 °C, 5% CO_2_, and humidity control), with a 10× objective lens over an 18-hr period. Control over opto-stimulation frequency was achieved by adjusting on/off exposure windows.

### Western Blotting

Cells grown were lysed with RIPA buffer containing Halt proteinase and phosphatase inhibitor cocktail (Thermo Fisher Scientific). Total protein concentration obtained through BCA protein assay (Pierce). Electrophoresis was performed using NuPAGE 4–12% Bis-Tris Gels, and transfer to PVDF membranes for Western Blotting was done in a tank blotting cell for 2 hours on ice. Membranes were incubated with primary antibody overnight at 4C, at the following dilutions: SMAD1 (Santa Cruz, 1:100) SMAD5 (Cell Signaling, 1:1000). Horseradish-peroxidase conjugated secondary antibodies (Sigma-Aldrich) were used at 1:100000 dilutions and incubated with the membranes for 1 hour at room temperature. Imaging SuperSignal West Dura ECL substrate (Pierce) in a Bio-Rad imager and quantification was performed using the Analyze Gels plugin from Fiji^60^.

### Immunocytochemistry

Cells were fixed in 4% paraformaldehyde in PBS for 10 minutes. After washing thoroughly with PBS, cells were permeabilized with 0.25 % Triton-X for 10 minutes on ice and blocked with 10% donkey serum at room temperature for 1 hour. Samples were afterwards incubated with primary antibodies overnight hours at 4 °C at the following dilutions: anti-tubulin β3 (1:1000; Sigma), anti-GFAP (1:1000; Abcam), pSMAD1/5 (1:400, Cell Signaling), mCherry (1:1000, Rockland), SMAD1 (1:100, Santa Cruz), Phalloidin-488 (Invitrogen, 1:100), Sox2 (Santa Cruz, 1:100). After several washes, samples were incubated with fluorescently-labeled secondary antibodies (Invitrogen) for 1 hour at room temperature. After two final washes, DAPI was added as a nuclear marker. Imaging of fixed-cell samples was performed on Perkin Elmer Opera Phenix confocal microscope, using 40× and 20× water-immersion objective lenses. All the image analysis and processing were performed with Fiji^60^.

### shRNA cloning

Oligos to build shRNA against SMAD1 were designed with Age I and Eco RI overhangs and obtained from Elim Biopharmaceuticals (shSMAD1.1-fw: CCGGGGTGCTCTATTGTGTACTATTCTCGAGTAGTACACAATAGAGCACCTTTTTTT; shSMAD1.1-rv: AATTCAAAAAAAGGTGCTCTATTGTGTACTACTCGAGAATAGTACACAATAGAGCAC C; shSMAD1.2-fw: CCGG*GTGCTCTATTGTGTACTATTTCTCGAGATAGTACACAATAGAGCACTT*TTTTT G; shSMAD1.2-rv: AATTCAAAAAAAGTGCTCTATTGTGTACTATCTCGAGAAATAGTACACAATAGAGCAC) and a LacZ control (Fw: CCGG*GGATCAGTCGCTGATTAAATTCTCGAGTTTAATCAGCGACTGATCCTT*TTTTT G; Rev: AATTCAAAAAAAGGATCAGTCGCTGATTAAA*CTCGAG*AATTTAATCAGCGACTGATC C). The oligos were annealed and cloned into the pLKO.1 hygro vector from Addgene (plasmid no. 24150).

### Viral Production and Cell Transduction

Lentiviral particles containing plKO-shRNA-SMAD1, LifeAct-mVenus or RhoA2G were packaged in human embryonic kidney (HEK) 293 T cells through polyethylenimine (PEI) transfection, and subsequently purified as previously described^64^. Retroviral vectors containing Cry2-mCherry-RhoA or Cry2-mCherry-Cdc42 developed before^26^ were packaged in HEK 293 T cells with Gag/pol and VSV-G through PEI transfection. NSCs were transduced at a multiplicity of infection (MOI) of 1, and selected using puromycin (1 μg/ml) or hygromycin (100 μg/ml) for 4 days. All vectors were verified by Sanger sequencing.

### Luciferase assay

NSCs were transduced with a lentiviral construct encoding a 7xTFP T cell factor/lymphoid enhancer factor (TCF/LEF) luciferase reporter, which represents β-catenin–TCF/LEF–based transcription. After the indicated treatments, cells were washed with PBS once, and lysed with lysis buffer (Promega). Suspensions were loaded into a white opaque 96-well plate, and Luciferase Assay Reagent (Promega) was added immediately before detection. Luminescence intensity was detected in a SpectraMax luminometer (Molecular Devices) and normalized to total protein concentration obtained through BCA protein assay (Pierce).

### Live-Cell Actin Imaging and Velocity Analysis

Actin cytoskeleton dynamics in live NSCs were studied using a LifeAct-mVenus biosensor^34^. Blue light exposure for opto-stimulation was accomplished by imaging in the 488-nm channel, one exposure per 2-5 s, with the 488-nm laser at 0.5% power using a confocal Zeiss LSM 880 NLO AxioExaminer with a 20× objective lens. Control over opto-stimulation frequency was achieved by adjusting on/off exposure windows. To image LifeAct-mVenus during the ‘off’ condition, 488-nm laser was used at 0.01% power, rendering enough signal-to-noise ratio to track the LifeAct probe without triggering Cry2 clustering. When pulses were longer than 15 minutes, LifeAct imaging was performed during the last 15 minutes of each ‘on’ or ‘off’ phase.

For velocity analysis of actin bundles, contrast on each image was first corrected using adaptive histogram equalization and image background was removed using a difference of gaussians filter in Fiji^60^. Peaks at least 20% above background in the resulting foreground image were then detected using non-local maximum suppression. Single-particle analysis of trajectory and velocity performed with the TrackMate plugin for Fiji^60^, was used to track subcellular dynamics of actin bundles (clusters of ∼500 nm in diameter).

LifeAct sensors fused to fluorescent proteins in their C-terminal, such as the LifeAct-mVenus fusion protein used in this study, have been shown to selectively interact with F-actin over G-actin^65^. To further determine whether actin fluctuations described here corresponded to F-actin or G-actin dynamics, cells expressing LifeAct-mVenus were fixed and stained for mVenus and Phalloidin (which specifically detects F-actin). The LifeAct/Phalloidin intensity ratio per cell was calculated with Fiji^60^. This ratio was nearly 1 within all cells analyzed and, in addition, it was was not significantly affected by the substrate stiffness (Supplementary Fig. 7). This indicates that the differences in actin fluctuations seen between these conditions represent differential F-actin bundles spatiotemporal dynamics.

### Statistical analysis

All the quantitative data are presented as mean ± SD, and the number of biological and technical replicates is indicated in the figure legends. One-way analysis of variance (ANOVA) followed by Tukey or Student’s *t* test for between-group differences were performed with Python. Kolmogorov-Smirnov test was used to evaluate statistical significance between distributions, and was performed with Python.

